# Nanopanel2 calls phased low-frequency variants in Nanopore panel sequencing data

**DOI:** 10.1101/2020.11.06.370858

**Authors:** Niko Popitsch, Sandra Preuner, Thomas Lion

## Abstract

Clinical decision making is increasingly guided by accurate and recurrent determination of presence and frequency of (somatic) variants and their haplotype through panel sequencing of disease-relevant genomic regions. Haplotype calling (phasing), however, is difficult and error prone unless variants are located on the same read which limits the ability of short-read sequencing to detect, e.g., co-occurrence of drug-resistance variants. Long-read panel sequencing enables direct phasing of amplicon variants besides having multiple other benefits, however, high error rates of current technologies prevented their applicability in the past. We have developed nanopanel2 (np2), a variant caller for Nanopore panel sequencing data. Np2 works directly on base-called FAST5 files and uses allele probability distributions and several other filters to robustly separate true from false positive calls. It effectively calls SNVs and INDELs with variant allele frequencies (VAF) as low as 1% and 5% respectively and produces only few low-frequency false-positive calls. Haplotype compositions are then determined by direct phasing. Np2 is the first somatic variant caller for Nanopore data, enabling accurate, fast (turnaround <48h) and cheap (sequencing costs ~10$/sample) diagnostic workflows.

## Main

Diagnosis and treatment of cancer patients is strongly guided by knowledge about the composition of disease-relevant (sub-)clones in patient samples. This requires technologies that can accurately determine the relative frequencies of haplotypes that inform about co-occurrence of variants on the same chromosome (compound mutations) which often play a central role in drug resistance^1^. For example, resistance to tyrosine kinase inhibitors including the third-generation drug ponatinib is caused by multiple mutations on the same allele in the tyrosine kinase domain (TKD) of BCR-ABL1 fusions in chronic myeloid leukemia (CML) patients^2–4^.

High-throughput panel sequencing has become a widespread tool for the rapid and cost-effective detection of such clinical biomarkers, among other reasons, due to its superior detection limit (LOD ~1% VAF) when compared to traditional methods (e.g., LOD of Sanger sequencing ~20% VAF). Nanopore long-read panel sequencing offers multiple benefits when compared to current short-read technologies: (i) direct phasing of variants within an amplicon, (ii) lower up-front and running costs, (iii) shorter interaction and turnaround times, (iv) accessibility of highly-repetitive genomic regions and (v) direct detection of structural variations. The relatively high error rates of Nanopore reads, however, made it difficult to call low-frequency variants in such datasets without producing large numbers of false-positive calls^5–9^.

### Nanopanel2

We have developed np2, a software that calls small somatic variants (SNVs and INDELs) from Nanopore panel sequencing data and iterates all identified haplotypes per sequenced amplicon. Briefly, np2 aligns base-calling probabilities from guppy to the specified reference (amplicon) sequence and then iterates these alignments column-wise (pileup) to call variants. Optional sample demultiplexing and alignment downsampling can be configured. Np2 calculates a number of statistics from aligned bases/INDELs per pileup and applies nine different algorithms to effectively filter false-positive calls (Sup. Fig. S1, Sup. Table S5). Our software then outputs VCF v4.2 and TSV files for downstream processing that contain all evaluated variant calls (including filtered ones) and associated core statistics. Finally, np2 calculates direct haplotypes for all non-filtered (PASS) variants and produces count tables and haplotype maps for visual interpretation.

### Read mapper selection

Long read alignment is non-trivial and error prone due to the high error rates in the used technology (a current review estimates ~87-98% accuracy^10^). We therefore evaluated the influence of different alignment algorithms on the overall performance of np2. Our software supports three different long-read alignment algorithms: minimap2 (mm2^11^), ngmlr (ngm^12^) and last^13^ and can be configured to align reads with any combination of these programs. If multiple aligners were used, np2 can calculate a (majority vote) consensus call set from all produced alignments. We measured per-mapper performance using data created from the Horizon OncoSpan reference standard (oncospan) and from a set of BCR-ABL1 TKD (abl1) panel sequencing datasets (Sup. Table S2, Sup. Fig. S3, Methods). Overall, mm2 showed the best performance with respect to false-discovery rates (FDR), and false-negative rates (FNR) while ngm and last resulted in higher numbers of false-negative (FN) and false-positive (FP) calls respectively (Fig. 1a, Sup. Fig. S4+S22). Mm2 datasets also showed the highest correlation between expected and observed VAF (Sup. Fig. S5). We therefore conducted our main analysis using mm2 alignments unless stated otherwise.

**Figure 1:**
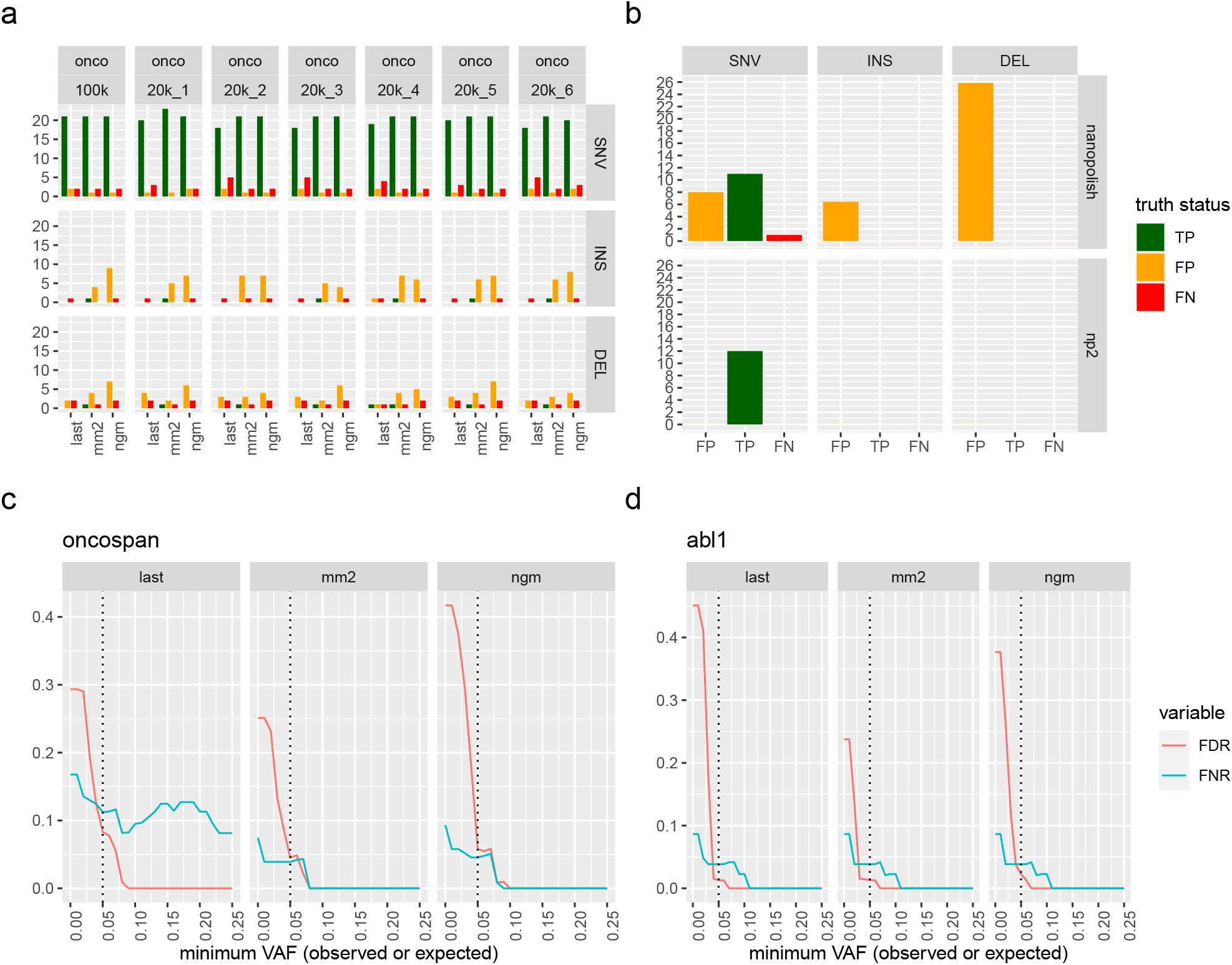
Np2 performance analysis. **a**, Number of true positive (TP), false positive (FP) and false negative (FN) variant calls per oncospan dataset, caller and variant type for all calls with an observed or expected (for FN) VAF > 5% (See Sup. Fig S12 for a corresponding plot without VAF threshold). The plot shows considerable numbers of FP INDEL calls, most of which, however, have low VAF (cf. Sup. Fig. S8). **b**, Mean numbers of TP, FP, FN calls for nanopolish (top panel) and nanopanel2 (bottom panel) in the 7 oncospan replicates. The evaluation was restricted to calls with expected/observed VAF>10% (Methods). Np2 consistently calls all 12 expected SNVs and produces no FP calls above this VAF threshold. Nanopolish misses one 25% VAF ALK SNV and produces a large number of false positive calls across variant types (SNV, INS, DEL). **c** and **d**: Mean false-discovery rate (FDR = FP / (FP + TP)) and false-negative rate (FNR = FN / (FN + TP)) per mapper and minimum VAF for all oncospan (**c**) and abl1 (**d**) SNV calls. The rapid drop of these measures with increasing minimum VAF confirms that most false calls have low VAFs. Minimap2 shows the best overall performance while last and ngmlr suffer from increased numbers of FN/FP calls respectively. Note, however, that FDR in **d** might still be inflated as we don’t have a complete ground truth for clinical datasets.

### Performance evaluation

We first evaluated np2 with seven downsampled oncospan alignments using variant calls from corresponding Illumina whole-exome sequencing (WES) data as ground truth (Fig. 1a). Overall, we observed 4 recurrent FN calls (Sup. Fig. S6, Sup. Table S6) and a reasonably small number of FP calls, most of which were low-frequency insertions. SNV FDR and FNR dropped quickly from 25% and 9% (1% minimum VAF) to zero (10% minimum VAF) with increasing VAF threshold (Fig. 1c). For abl1 data, we observed similar numbers with FDR and FNR rapidly dropping from ~24% and 9% to zero when reaching a minimum VAF threshold of 11% (Fig. 1d, Sup. Fig. S7). Moreover, FDR of clinical abl1 data is possibly inflated as we were unable to validate all low-frequency variants due to the detection limits of the used validation methods. Due to the low number of validated insertions and deletions in our datasets we could not conduct the same performance analysis for INDELs. Np2 did, however, call 13 out of 21 (TPR 62%) INDELs (2 deletions and one insertion with mean expected VAF 5.3%) over all oncospan replicates and did not produce any unfiltered high VAF INDEL calls (Sup. Fig. S8+S9). Together, these results confirm that np2 can call low-frequency SNVs and INDELs with acceptable false discovery rates and reliably calls variants with VAF above 10%. Further investigating the produced FPs we could observe a high degree of recurrence (Sup. Fig. S10) which indicates a systematic cause and offers possibilities for further optimisations, e.g., by filtering variant c alls w ith high recurrence across multiple independent samples.

### Filtering stats

We then investigated which of np2’s filters are most effective for suppressing FP calls by analysing the filter subsets assigned to >200k true-negative (TN) calls in 40 samples (Sup. Fig. S2). For SNVs, we found our novel allele quality filter (AQ1, Methods) to be the single most effective filter preventing 375 false-positives that would not have been detected by any other filter. Briefly, AQ1 is based on recalculating per-base qualities in an alignment column by incorporating the distributions of all allele probabilities as provided by guppy (Methods). For INDELs, allele frequency was the most effective single filter which is a direct result of using respective VAF thresholds in our evaluation runs. Besides this, filtering for strand bias (SB) and homopolymer runs (HP) seemed particularly effective for all variant types (Fig. 2a).

**Figure 2:**
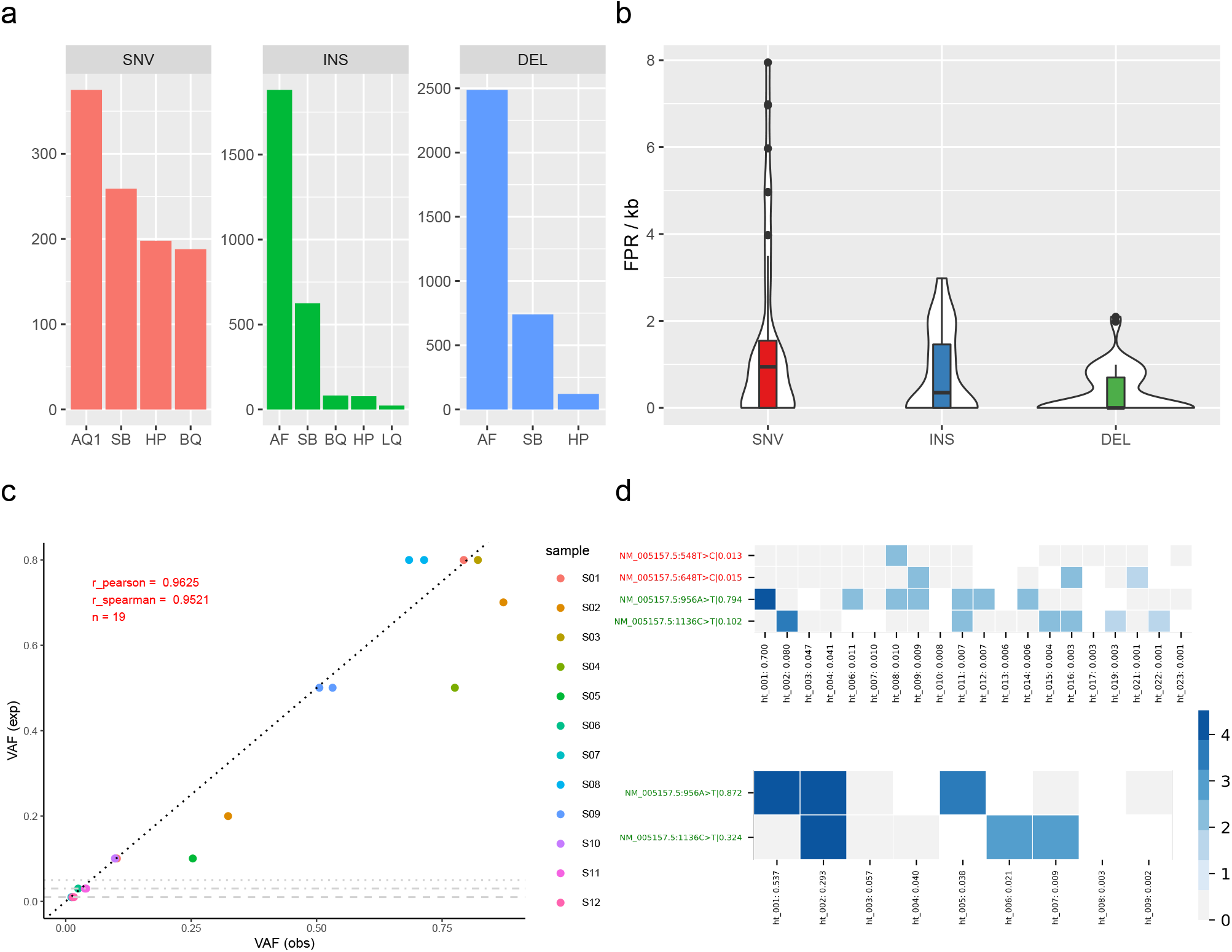
**a**, TN calls per variant type across 40 samples that were filtered exclusively by one single np2 filter. The plots reveal that our allele quality filter (AQ1) is the single most effective filter for suppressing FP SNV calls followed by strand bias (SB) and homopolymer (HP) filters. INDEL FPs are predominately filtered due to low allele frequency (AF), SB and HP filters. A more detailed analysis is provided in Sup. Fig S5. **b**, False-positive rates (FPR) per kb amplicon for all evaluated datasets. Overall we found medium FPR values of 0.95 and 0.7 for SNVs and INDELs. FPR per individual dataset is shown in Sup. Fig. S11. **c**, Strong correlation between expected (exp) and observed (obs) VAF for 18 (single and compound) benchmark variants of abl1_min_1, particularly for low-frequency variants. Compound variants have equal expected and similar observed VAF (e.g., S08; all 12 samples are described in Sup. Table S7). One variant with an expected VAF of 1% was not called by np2 and is not shown, horizontal grey lines show calling thresholds for deletions, insertions and SNVs respectively. **d**, Exemplary haplotype maps as plotted by np2 for two benchmark samples containing the same mutations (Y axis, green labels. Red labels denote low-frequency false positive calls; raw VAF as determined by np2 after the pipe symbol). The X axes list found haplotypes sorted by their relative fraction (number after the colon). Blue and grey squares denote present/absent variations, white squares indicate that the respective reads did not span the respective positions or had a deletion at this position. Colour intensity corresponds to log(#reads). In the upper sample, both mutations were introduced using different plasmids. Targeted VAFs were 80% E255V (ht_001), 10% T315I (ht_002) and 10% wt (ht_003+4) respectively. The lower sample contained a mix of 70% single E255V (ht_001), 20% compound E255V+T315I (ht_002) and 10% wt (ht_003+4). The haplotype maps confirm the respective linkage status of the benchmark variants although we find a low-frequency ht in the upper map carrying both mutations (ht_011; freq=0.7%), possibly resulting from sequencing errors.

### Nanopolish comparison

As np2 is the first available somatic variant caller for Nanopore data we chose to compare its performance to Nanopolish^14^, a tool that has repeatedly been used for germline variant calling (Fig. 1b). This evaluation was restricted to oncospan calls with VAF>10% due to technical limitations of Nanopolish (see Supplement). Np2 called all expected variants for this dataset and did not produce any FP calls above this VAF threshold. In contrast, Nanopolish missed one variant in the ALK gene and produced a large number of false positive SNV and INDEL calls. While these results are reasonable because Nanopolish was not developed as a somatic variant caller they underline np2’s ability to efficiently suppress high-frequency FPs. To further demonstrate this, we calculated false-positive rates per kilobase amplicon (FPR) across all datasets with np2’s default VAF thresholds of 1% for SNVs, 3% for insertions and 5% for deletions respectively and found median FPR values of 0.95 and 0.7 respectively for SNVs and INDELs (Fig. 2b, Sup. Fig. S11; note that the lower FPR for INDELs results from the more stringent AF filtering) which correspond to one FP SNV/INDEL call per 1053/1429 amplicon bases respectively.

### Reproducibility and detection limits

To measure how accurately np2 can reproduce expected VAFs from a plasmid dilution series, we created two benchmark datasets with known concentrations of variant-carrying plasmids (Methods). We found a high correlation (r pearson 0.96 and 0.98 respectively, Fig. 2c, Sup. Fig. S5) between expected and observed VAF values. Furthermore, we observed very similar reported VAFs for pairs of variants located on the same plasmid. Np2 called 28 of 31 variants (90%) down to a VAF of 1%. All three FN SNV calls had an expected VAF of 1% and when considering only such low-frequency variants, np2 called 4/7 (FNR=0.43).

We were interested in the minimum required read depth for np2 and randomly downsampled the data from a MinION flow-cell (abl1 m in 3) to sets of 1k, 5k, 10k and 25k reads per sample respectively. We then compared the resulting np2 call sets and found high correlation despite the relatively large difference in read numbers (Sup. Fig. S13). TPR compared perfectly between the downsamples (Sup. Fig. S14). Encouraged by these results, we sequenced one benchmark sample and 8 multiplexed clinical samples on two ‘Flongle’ flow-cells to evaluate the suitability of these smaller flow-cells for clinical applications. Across these 9 samples we called all 10 expected variants and again obtained only few low-frequency FPs (Sup. Fig. S18). Additionally, we observed good variant calling performance (no FN and only 3 FP over 7 replicates) on a lowly-covered oncospan amplicon (KIT 1) with a median coverage of only 330 reads in the 20k samples (Sup Table S2). Together, this suggests that np2 does not require high read depth to accurately call somatic variants and indicates its possible application in targeted Nanopore sequencing^15, 16^.

### Conclusions

Nanopanel2 is the first haplotype aware somatic variant caller for Nanopore panel sequencing data and our analysis shows that np2 accurately calls variants above 10% VAF and reaches good performance for low-frequency variants down to its currently recommended thresholds of 1% VAF for SNVs and 5% VAF for INDELs. Reported VAFs showed very high correlation to the expected values. Furthermore, np2 provides information on the distribution of subclones in a sample by directly calling haplotypes from (amplicon-spanning) long reads. Together, this makes it a viable alternative to current amplicon sequencing methods for clinical and research applications.

Despite the promising results of our evaluation using benchmark and clinical samples, more work is required to improve np2’s calling accuracy. One particular problem is the accurate calling in and around homopolymer runs, a known problem of Nanopore’s current sequencing technology^6^. Although our hp filter effectively removes most FPs in such regions, this also results in FN calls and for the time being, we recommend users to revalidate calls that were filtered exclusively by the HP filter (see Methods). We do, however, expect this problem to be mitigated in the future due to recent improvements in nanopore chemistry^17^.

Nanopore’s technology enables fast and cost-efficient on-demand sequencing which is particularly attractive for small diagnostic labs that cannot wait until larger (short-read) flow-cells have been filled with multiplexed patient samples. Cost efficiency is also key for clinical management that includes early detection and continuous monitoring of disease relevant (resistance) clones^8^. Here, we have showcased a clinical diagnostics workflow for CML patients that is cheap (e.g., 8 multiplexed samples on a single 90$ Flongle flow-cell with ~$50 library preparation costs) and fast (library prep 2-3h, sequencing: 12-24h, bioinformatics ~14h).

Besides clinical applications, np2 has other obvious usage scenarios including metagenomic and full-length virus sequencing^18–20^. Our experiments with low-coverage amplicons/downsamples furthermore indicate that np2’s variant filtering approach might be adaptable to target-enriched^15, 16^ or even whole-genome nanopore data^21^. We furthermore speculate that our allele quality filtering method could be combined with current approaches for read polishing and germline variant calling^22–24^.

## Methods

### Oncospan data

We obtained Nanopore sequencing data for 9 amplicons from the cell-line-derived Horizon OncoSpan gDNA reference standard (https://horizondiscovery.com/, cf. Sup. Table S3) from Oxford Nanopore Technologies Ltd. The oncospan DNA samples were sequenced on a single MinION flow-cell resulting in overall 13.5 Mio reads. From these data we created six samples with 20k reads each and one sample with 100k reads by random subsampling (Sup. Table S2). Additionally, we obtained deep Illumina WES data (500X coverage; from provider) for these amplicons, called (low-frequency) variants using the pisces^25^ variant caller (v5.2.9.122, command lines are provided in Supplement) and manually inspected variant calls in a genome browser. We then considered all regions with a minimum coverage of 10X in Nanopore and Illumina data and measured np2’s performance considering unfiltered pisces calls as ground truth. Note that a subset of these variants was validated by droplet digital PCR (ddPCR) in the original oncospan data.

### BCR-ABL1 data

We further evaluated np2 using manually created in-vitro benchmark datasets as well as several clinical datasets in a BCR-ABL1 background with a single amplicon covering the TKD of the BCR-ABL1 fusion gene. First, we created a dilution series (‘abl1 min 1’, Sup. Table S7) using Thermofisher Topo T4 cloned plasmids containing major BCR-ABL1 constructs (wild-type, T315I single mutation, E255V single mutation and T315I+E255V compound mutations). We used either one plasmid containing both mutations (compound) or two plasmids containing one mutation each which allowed us to benchmark np2’s ability to call haplotypes. Plasmids were calibrated to a BCR-ABL1 copy number of 1E+4 by RT-PCR^26^. T315I single- and T315I+E255V compound plasmids were then mixed into the BCR-ABL1 WT background (VAF 80%, 50%,10%, 5%, 3%, 1%).

We also created two replicates of a dilution series (Supl. Table S8, ‘abl1 min 2’/dilution) containing two compound variants at different concentrations (10%, 3%, 1%) to learn about np2’s detection limit and about how well it could reproduce expected VAFs. Additionally, we evaluated np2 with different combinations of clinically relevant variants from an international ring trial comprising cDNA samples derived from BCR-ABL positive cell lines with a BCR-ABL transcript copy number of about 10% (‘abl1 min 2’/ringtrial).

To learn about the influence of read depth on np2’s performance, we sequenced a MinION flow-cell with 9 clinical samples (‘abl1 min 3, Sup. Table S9) and randomly downsampled the alignments to 1k, 5k, 10k and 25k reads per sample. Additionally, we sequenced one benchmark sample (plasmid mix, ‘abl1 flo 1’) and 8 multiplexed clinical samples (‘abl1 flo 2’) on two ‘Flongles’ (Sup. Table S10) to evaluate np2’s performance on these smaller flow-cells.

The truth status of variants in clinical samples was determined/validated using various orthogonal methods including Sanger sequencing^27^, ligation-dependent PCR^28^, MiSeq panel sequencing^29^ and Pyrosequencing^30^. Notably, we do not have a complete ground truth for clinical samples due to the different detection limits of the used validation techniques (e.g., 20% for Sanger sequencing). Consequently, false-positive rates for the respective datasets might be inflated as we don’t know whether low-frequency variants called by np2 are in fact true variants.

### Nanopanel2 preprocessing

Np2 takes guppy (v3.6.1) base-called FAST5 files and a JSON configuration file (we used np2’s default configuration for all evaluation runs) as input and optionally starts by demultiplexing reads using porechop (v0.2.4, ^22^) if configured. It then extracts FASTQ files and guppy probability values from the FAST5 files. In this step, np2 can optionally split reads that exceed a configurable length as we observed considerable numbers of chimeric reads in the oncospan data (possibly artefacts from the PCR reaction). Reads are then aligned to the reference sequence using the configured long-read mappers and BAM files are annotated with extracted base-calling probability values for efficient downstream processing. Optionally, np2 can randomly downsample the final alignments, e.g., for faster processing or debugging purposes.

### Variant calling

Np2 iterates alignments column-wise and, for each position, considers all alternate alleles (alt) with a configurable minimum number of supporting reads per pileup as potential sequence variants. It then interrogates guppy probability distributions for each alt base-call in the pileup and filters reads that show increased probabilities for any different allele (including the reference allele) among those. For this, np2 first iterates the maximum ‘flip-flop’ probability for each possible non-alt allele and then tests for equal distribution of these values using a *χ*^2^ test, rejecting base-calls with skewed distributions (p≤0.05) that we found to be error prone. In initial experiments, we observed that false positive SNV calls were often correlated with high overall ref base probabilities in the pileup, demonstrating that the basecaller was unsure about its decision to call the alt allele. We then use this algorithm to calculate corrected alt-allele counts and frequencies and filter potential SNV calls based on configurable thresholds for these values (filter ‘AQ1’). Np2 furthermore filters calls with large fractions of rejected alt-base calls (AQ2). S up. F ig. S19a+b show, for example, the effect of this correction algorithm around homopolymer runs.

Strand bias, meaning that the alt allele is disproportionally found on reads of a particular strand, is a strong indicator of artefacts in NGS data. We have implemented a strand-bias (SB) filter for SNV calls based on testing contingency tables created from plus and minus strand counts of ref and alt reads with a *χ*^2^ test with Yates correction (p≤0.05). Prior to testing, contingency tables are corrected for the flow-cell/sample specific strand ratio calculated from all sequenced reads (cf. Sup. Fig. S23). Furthermore, the SB filter is applied only if alt allele carrying reads show a strong imbalance of supporting plus and minus strand reads (*aa_skew* = |*log*2(#*minus/*#*plus*)| > *threshold*). INDELs are filtered only for the latter criteria.

Nanopore’s current technology experiences problems in accurately calling the length of homopolymer (hp) runs (Rang2018,Orsini2018), which consequently resulted in high numbers of false-positive calls at the hp run borders before filtering (data not shown). Sup. Fig. S19d plots histograms of hp runs with minimum length 3 in the considered amplicons and shows that overall around 23% of all bases reside in such regions. To mitigate this problem, we have implemented a homopolymer filter (HP) that calculates hp lengths up- and downstream of the current pileup and filters calls if those are exceeding a given threshold (default 3 and 2 for up/downstream hp runs) and if one of these hp runs consists of the alt allele (indicating misalignment). For INDELs, we add the lengths of up- and downstream alt-allele hp runs and filter if this sum exceeds a threshold (default: 3). Although this strategy effectively reduces FP calls, it comes at the cost that some calls at hp borders will always be filtered despite their other quality measures which may result in FNs. One example of this is the oncospan FN deletion MET 2:687GT>G (Sup. Fig. S21, Sup. Table S6). Another, clinically relevant, example is the ABL1 variant E255K, a G>A SNV at the end of a 4-base hp G run that is followed by an A in the reference sequence. For this reason, we recommend that users take a close look at calls that were filtered only by our HP filter and revalidate the respective calls using orthogonal methods where possible. In our datasets (40 samples) we found 398 such calls (an average of 5 SNVs and 5 INDELs per sample) with a high degree of recurrence (99 distinct calls). Only one (0.25%) of these calls was a true negative and the median VAF of calls was low: 2% (SNV), 4% (INS) and 8% (DEL) respectively (Sup. Fig. S20).

Finally, we calculate a Phred-scaled quality score [0; 100] based on different pileup statistics: for SNVs, this score is min(100, −*log*10(*non_alt_prob*)) where *non_alt_prob* is the mean of all normalised non-alternate allele guppy probabilities. For insertions and deletions it is based on hp length, *aa_skew* and average base quality (see Supplement for details). Variants are filtered as ‘low quality’ (LQ) based on configurable, variant type specific thresholds. Further threshold-based filters include low (raw) allele frequencies (AF), low read depth (DP), low mean base qualities (BQ) and high strand-imbalance (SI) that is based on the ratio of all covering plus and minus strand reads.

### Haplotype maps

Np2 will consider all unfiltered variant calls for calculating haplotype maps that may inform users about the distribution of sub-clones in a sample. First, we iterate all reads and check for presence of the respective alt/ref alleles. We then summarise this data per haplotype and draw a heatmap for visual inspection (Fig. 2d, Sup. Fig. S15-S17). For this heatmap, we filter the haplotypes by the following criteria: (i) minimum fraction among all considered reads >0.1%, strand skew *log*2(#*plus_strand_reads/*#*minus_strand_reads*) < 2, (iii) minimum number of supportive reads>10, (iv) a maximum of 25 haplotypes is shown. Blue squares in this heatmap indicate presence of the respective variant and colour intensity corresponds to *log*10(#*reads*). Grey squares correspond to the reference allele, white squares indicate non (most-frequent) alt or ref alleles or missing coverage of the respective reads.

### Performance evaluation

Using the respective truth sets we count FP, TP, TN and TP calls per sample and calculate false-discovery rates 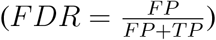, false-negative rates 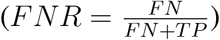,true-positive rates 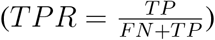 and false-positive rates 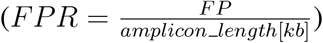 where appropriate.

## Data availability

The data for this study have been deposited at zenodo.org under DOIs accession numbers 4110691 and 4110698.

## Acknowledgements

The authors thank Philipp Rescheneder and Phillip James (Oxford Nanopore Technologies Ltd.) for providing the oncospan datasets and helping with data interpretation and Thomas Ernst (University Jena, Germany) for the permission to test samples with quantified mutation loads derived from the EUTOS international control round for deep sequencing analysis of BCR-ABL mutations in CML.

## Author information

N.P., T.L. and S.P. conceptualised the project and designed the experiments. N.P. designed and implemented the software and conducted all *in-silico* experiments. S.P. prepared and sequenced all BCR-ABL1 datasets. All authors were involved in interpretation of the data, discussed the results and commented on the manuscript. N.P. wrote the manuscript with contributions from all authors.

