## Supplemental Data for "Nanopanel2 calls phased low-frequency variants in Nanopore panel sequencing data"

Supplementary Information for
*Nanopanel2 calls phased low-frequency variants in Nanopore panel sequencing data*

Niko Popitsch[1,2,*], Sandra Preuner[2] & Thomas Lion[2,3]

[1]Institute of Molecular Biotechnology of the Austrian Academy of Sciences (IMBA)
Vienna Biocenter (VBC), Dr. Bohrgasse 3, 1030 Vienna, Austria.
[2]Children's Cancer Research Institute, Zimmermannplatz 10, 1090 Vienna, Austria.
[3]Department of Pediatrics, Medical University of Vienna,
Währinger Gürtel 18-20, 1090 Vienna, Austria.
∗Corresponding author

**II Supplementary Tables** **6**

**III Supplementary Figures** **12**

#### Introduction

The following document contains supplementary information and figures for the paper 'Nanopanel2 calls phased low-frequency variants in Nanopore panel sequencing data'. It is split into three parts. Part I contains various text sections providing additional details about the conducted evaluation and nanopanel's algorithms. Parts II + III contain supplementary tables and supplementary figures showing additional data/analyses as referenced throughout the main manuscript.

### Part I

### Supplementary Text

### 1 Pisces variant calling in short-read WES data

The following commandline was used to call somatic variants in the oncospan WES dataset with pisces v5.2.9.122

```
pisces -bam bam_file -CallMNVs false -g fasta_file -gVCF false -OutFolder out_dir -RMxNFilter 5,9,0.35
          -MinVF 0.0005 -SSFilter false -MinVQ 0 -MinDepth 5
```

Unfiltered calls were considered as truth set for the oncospan dataset.

### 2 Long read mapping

##### Reference genomes

ABL1 data was mapped against $NM\_005157.5$, the respective FASTA file can be downloaded from NCBI. The reference sequence for the sequenced Oncospan amplicons was deposited at XXX. The following commandlines were used to map reads using the respective long read mapper:

##### last v1042

```
last lastal -Q1 last_db fastq_file > tmp_file
last last-split < tmp_file > maf_file
last maf-convert SAM maf_file > sam_file
```

We then added missing headers (@HD, @PG, @SQ) to the resulting SAM files, converted them to BAM format and extracted all alignments with maximum alignment score (AS tag) to ensure a single (primary) alignment per read name. Finally, resulting BAM files were sorted and indexed.

##### ngmlr v0.2.7

```
ngmlr -r fasta_file -q fastq_file -x ont -t threads -o sam_file --no-smallinv --no-lowqualitysplit \
      -k 10 --match 3 --mismatch -3 --bin-size 2 --kmer-skip 1
```

Resulting SAM files were converted to BAM, sorted and indexed.

##### minimap2 v2.17-r941

```
minimap2 -ax map-ont -t threads fasta_file fastq_file -o sam_file
```

Resulting SAM files were converted to BAM, sorted and indexed.

#### 3 Nanopolish evaluation

We compared np2 to Nanopolish v0.13.2. This evaluation was restricted to oncospan calls with expected/observed VAF>10% due to technical limitations of Nanopolish. We were unable to produce low-VAF calls with Nanopolish because it does not call segments with >200 call candidates (built-in, hard threshold) which we easily exceeded using e.g., a 5% VAF threshold and because Nanopolish does not work with ploidy>2 as it was not developed as a somatic variant caller. The following commandline was used to create the Nanopolish call sets:

```
nanopolish variants -o out_file \
                    -t threads \
                    -r fastq_file \
                    -b bam_file \
                    -g fasta_file \
                    --min-candidate-frequency 0.1 \
                    -w roi_interval \
                    --ploidy 2 \
                    --calculate-all-support \
                    --max-haplotypes 1000000 \
                    -v
```

### Part II

### Supplementary Tables

| Variant type | Formula | Rationale |
|---|---|---|
| SNV | $quality = min(100, -log10(non\_alt\_prob))$ | pileups with high non-alt allele probabilities are unreliable |
| INS | $data = [1 - min(hp\_len, 5)/5]$. | long HP => low score |
| | $data+ = [0]$ if $seq\_eq\_ins$ else $[1]$ | ref seq equals inserted bases => low score |
| | $data+ = [1 - min(5, abs(aa\_skew))/5]$ | high aaSkew => low score |
| | $data+ = [min(30, avg\_base\_qual)/30]$ | high avg base quality => high score |
| | $quality = min(100, -10 * log10(1 - mean(data)))$ | call quality phred score calculation |
| DEL | $data = [1 - min(hp\_len, 5)/5]$ | long HP => low score |
| | $data+ = [1 - min(5, abs(aa\_skew))/5]$ | high aaSkew => low score |
| | $data+ = [min(30, avg\_base\_qual)/30]$ | high avg base quality => high score |
| | $quality = min(100, -10 * log10(1 - mean(data)))$ | call quality phred score calculation |

Table S1: Nanopanel2's call quality (Phred score) calculation per variant type in pseudo code.

| Panel | Dataset | Amplicons | Flowcell | Samples | Multiplexed | Description |
|---|---|---|---|---|---|---|
| ABL1_NM_005157.5 | abl1_min_1 | 1 | MinION | 12 | yes | benchmark variants (single and compound) |
| ABL1_NM_005157.5 | abl1_min_2 | 1 | MinION | 12 | yes | Dilution series (barcodes 1-6) and ring trial variants (barcodes 7-12) |
| ABL1_NM_005157.5 | abl1_min_3 | 1 | MinION | 9 | yes | clinical samples (downsampled) |
| ABL1_NM_005157.5 | abl1_flo_1 | 1 | Flongle | 1 | no | benchmark variants |
| ABL1_NM_005157.5 | abl1_flo_2 | 1 | Flongle | 8 | yes | clinical samples |
| Oncospan | onco_20k_1 | 9 | MinION | 1 | no | benchmark variants |
| Oncospan | onco_20k_2 | 9 | MinION | 1 | no | benchmark variants |
| Oncospan | onco_20k_3 | 9 | MinION | 1 | no | benchmark variants |
| Oncospan | onco_20k_4 | 9 | MinION | 1 | no | benchmark variants |
| Oncospan | onco_20k_5 | 9 | MinION | 1 | no | benchmark variants |
| Oncospan | onco_20k_6 | 9 | MinION | 1 | no | benchmark variants |
| Oncospan | onco_100k_1 | 9 | MinION | 1 | no | benchmark variants |

Table S2: Datasets used in this study

| Panel | Amplicon | Gene | Length | Mean coverage in 20k oncospan samples |
|---|---|---|---|---|
| ABL1_NM_005157.5 | NM_005157.5 | ABL1 | 5,596 | - |
| Oncospan | ALK | ALK | 1,100 | 20,950 |
| Oncospan | EGFR_1 | EGFR | 1,010 | 21,915 |
| Oncospan | EGFR_2 | EGFR | 990 | 21,989 |
| Oncospan | EGFR_3 | EGFR | 1,070 | 21,311 |
| Oncospan | KIT_1 | KIT | 1,030 | 330 |
| Oncospan | KIT_2 | KIT | 890 | 22,215 |
| Oncospan | MET_1 | MET | 990 | 21,971 |
| Oncospan | MET_2 | MET | 1,040 | 21,250 |
| Oncospan | PIK3CA | PIK3CA | 1,020 | 21,143 |

Table S3: Sequenced amplicons

| Dataset | Flowcell | Nanopore Version | Mean read length [bp] | Total Reads | Passed Reads | Failed Reads | Approx Runtime |
|---|---|---|---|---|---|---|---|
| oncospan | MinION | R9.4.1 | 1,079 | 13,572,202 | 11,691,353 | 1,880,849 | 48h |
| abl1_min_1 | MinION | R9.4.1 | 1,732 | 887,247 | 694,525 | 192,722 | 24h |
| abl1_min_2 | MinION | R9.4.1 | 1,606 | 704,709 | 535,271 | 169,438 | 24h |
| abl1_min_3 | MinION | R9.4.1 | 1,100 | 4,702,087 | 3,688,465 | 1,013,622 | 24h |
| abl1_flo_1 | Flongle | R9.4.1 | 1,540 | 237,298 | 144,590 | 92,708 | 24h |
| abl1_flo_2 | Flongle | R9.4.1 | 1,505 | 375,610 | 261,154 | 114,456 | 24h |

Table S4: Core sequencing statistics per dataset/flowcell

| ID | Name | Description |
|---|---|---|
| AF | Low allele frequency | Raw allele frequency below configured thresholds [SNV: 0.01, DEL: 0.05, INS: 0.03] |
| DP | Low depth | Raw read depth below configured thresholds [SNV: 10, DEL: 100, INS: 100] |
| BQ | Low base quality | Mean base quality below configured threshold [12] |
| SB | Strand bias | Strand bias ($p - value < 0.05$ and $\|aa\_skew\| > configured\_treshold$) [SNV: 1.0, DEL: 0.2, INS: 0.5] |
| AQ1 | Low allele quality filter1 | SNV only: Corrected allele frequency below configured thresholds [0.01] or corrected allele-count below threshold [10] |
| AQ2 | Low allele quality filter2 | SNV only: $Corrected\_allele\_count/raw\_allele\_count < configured\_treshold$ [0.3] |
| HP | Homopolymer filter | Homopolymer filter, see main manuscript for details |
| LQ | Low call quality | Call quality below configured threshold [SNV: 10, DEL: 3, INS: 3], see below for details |
| SI | Strong strand imbalance | Very high/low fraction of plus-strand reads [$< 0.2$ or $> 0.8$] |

Table S5: Short description of nanopanel2 filters and default thresholds (square brackets). See main manuscript for a more detailed description of AQ1, SB and HP.

| Chr | Pos | Ref | Alt | Type | VAF_exp | VAF_obs | Description |
|---|---|---|---|---|---|---|---|
| PIK3CA | 551 | G | A | SNV | 0.08182 | 0.008828432 | Filtered due to low AF and other filters in 6/7 subsamples; the VAF was approximately 10X underestimated (VAF_obs ∼0.9% vs VAF_exp 8%) |
| MET_2 | 687 | GT | G | DEL | 0.05245 | 0.13694436 | Filtered with strand-bias, homopolymer and low call quality filters in all 7 oncospan datasets. The VAF was ∼3X overestimated (VAF_obs ∼14% vs VAF_exp 5%). An IGV screenshot showing the short-read WES alignment at this position is shown in Sup. Fig S21 |
| KIT_2 | 501 | A | T | SNV | 0.07359 | NA | Not called in 6/7 subsamples due to too low allele fraction (e.g., in replicate 2: 144/21737 reads) |
| ALK | 135 | C | CATTG | INS | 0.05469 | NA | Not called in 1/7 samples. Instead, a 'C/CGATG' insertion was called in this sample (replicate 2). |

Table S6: Description of false negative calls in the oncosopan evaluation datasets. See Sup. Fig. S6 for additional data for these FNs.

| Sample | Description |
|---|---|
| S01 | E255V 80%, T315I 10% |
| S02 | E255V 70%, E255V+T315I 20% |
| S03 | T315I 80% |
| S04 | T315I 50% |
| S05 | T315I 10% |
| S06 | T315I 3% |
| S07 | T315I 1% |
| S08 | T315I + E255V 80% |
| S09 | T315I + E255V 50% |
| S10 | T315I + E255V 10% |
| S11 | T315I + E255V 3% |
| S12 | T315I + E255V 1% |

Table S7: Compound (A+B) and single (A, B) ABL1 mutations on flowcell 'abl1_min_1'.

| Sample | Description | Category |
|---|---|---|
| S01 | T315I+E255V 10% | Dilution series |
| S02 | T315I+E255V 10% | Dilution series |
| S03 | T315I+E255V 3% | Dilution series |
| S04 | T315I+E255V 3% | Dilution series |
| S05 | T315I+E255V 1% | Dilution series |
| S06 | T315I+E255V 1% | Dilution series |
| S07 | M244V 30%, T315I 30%, E282K 20% | Ring trial |
| S08 | M244V 5%, M351T 20% | Ring trial |
| S09 | Y253F 30% | Ring trial |
| S10 | H396P 20% | Ring trial |
| S11 | F311L 5% | Ring trial |
| S12 | L387M 10% | Ring trial |

Table S8: Compound (A+B) and single (A, B) ABL1 mutations on flowcell 'abl1_min_2'.

| Sample | Description | Patient | Category |
|---|---|---|---|
| S01 | N/A | A | minor BCR-ABL |
| S02 | Y253H 83% | A | minor BCR-ABL |
| S03 | Y253H 83%, F317L 7% | A | minor BCR-ABL |
| S04 | T315I 40%, Y253H 82%, F317L 12% | A | minor BCR-ABL |
| S05 | T315I 18%, Y253H 27%, E255V 58% | A | minor BCR-ABL |
| S06 | E255V 100% | A | minor BCR-ABL |
| S07 | T315I 10% | B | major BCR-ABL |
| S08 | N/A | B | major BCR-ABL |
| S09 | T315I 20%, E255K 22% | B | major BCR-ABL |

Table S9: Clinical sample mutations on flowcell 'abl1_min_3' as validated by Sanger sequencing

| Flongle | Sample | Description |
|---|---|---|
| abl1_flo_1 | S01 | T315I 50% |
| abl1_flo_2 | S01 | A269A 100% |
| abl1_flo_2 | S02 | M244V 100% |
| abl1_flo_2 | S03 | H396R 19% |
| abl1_flo_2 | S04 | E255V 100% |
| abl1_flo_2 | S05 | T315I 100% |
| abl1_flo_2 | S06 | E459K 100% |
| abl1_flo_2 | S07 | G250E 51% |
| abl1_flo_2 | S08 | E255V 100%, T315I 100% |

Table S10: Clinical sample mutations on flongles 'abl1_flo_1' and 'abl1_flo_2' as validated by Sanger sequencing

### Part III

### Supplementary Figures

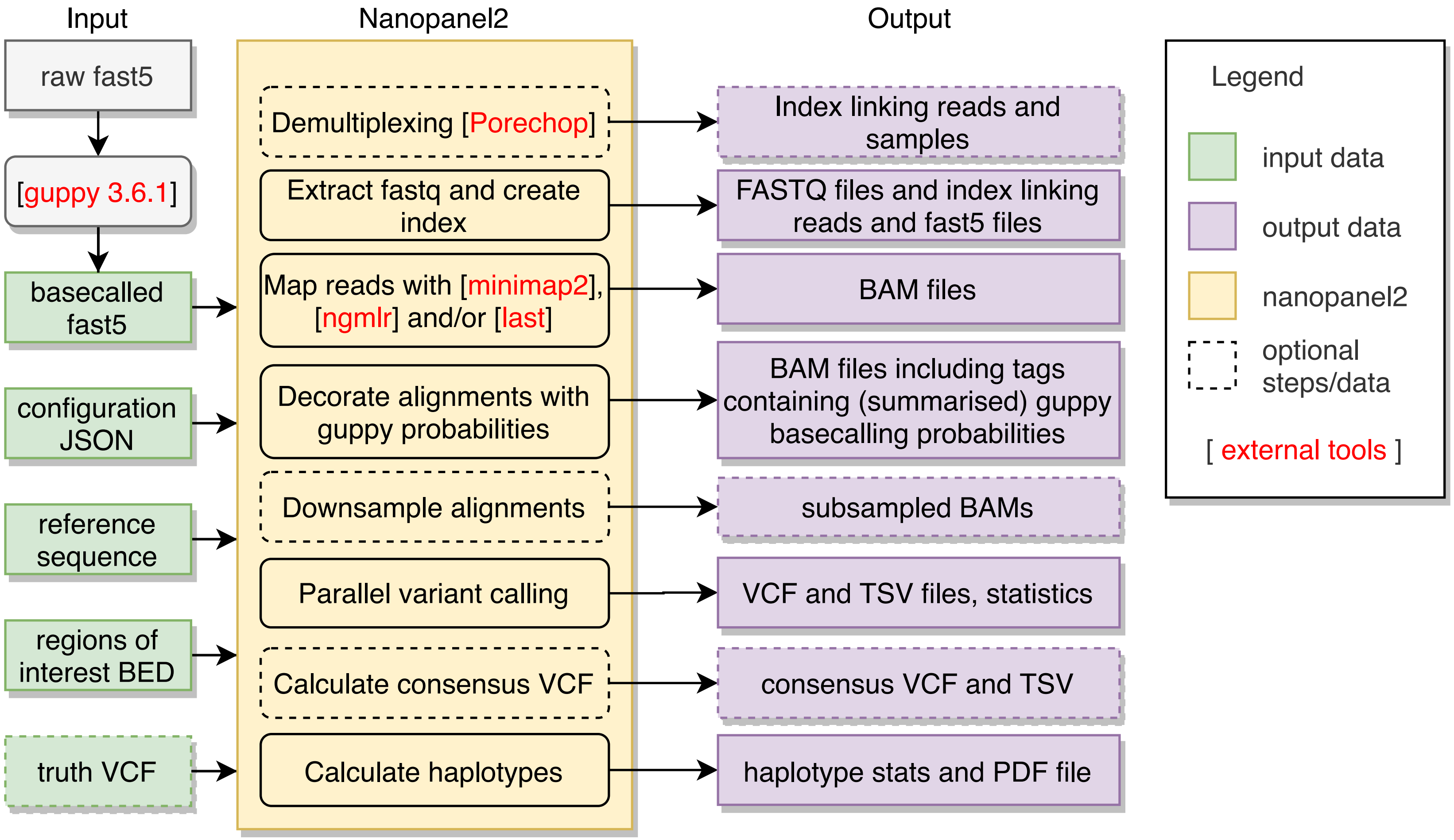

Figure S1: Block diagram showing the core components of Nanopanel2 (np2). Briefly, Nanopore reads are demultiplexed (optional), read bases and guppy basecalling probabilities are aligned to a given amplicon reference sequence and variants and haplotypes are called. See Methods for a detailed description of the individual steps. Np2 is implemented in python3.7 and makes use of external tools as indicated (including samtools and htslib; not shown).

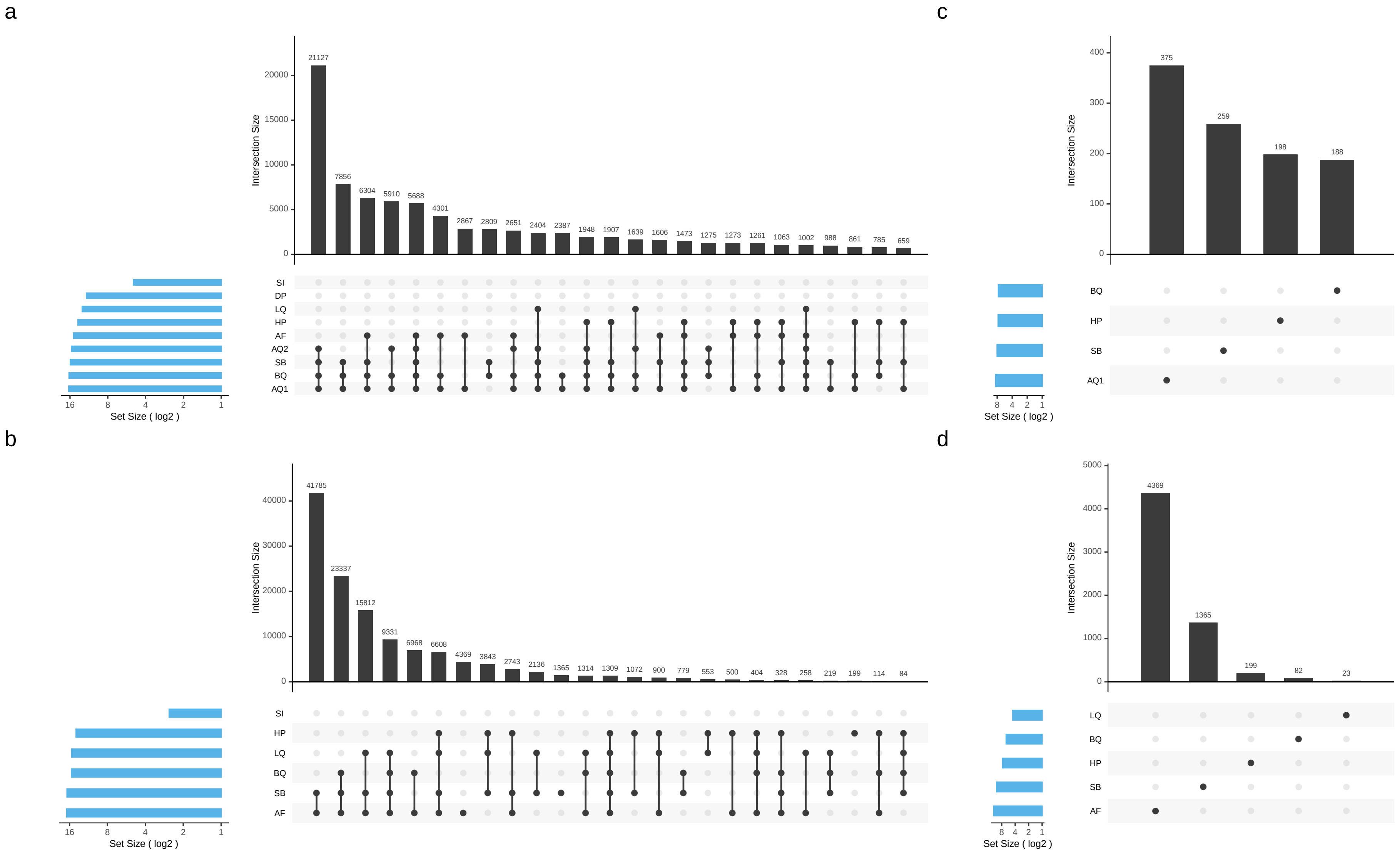

Figure S2: Distribution (UpSet plots, Lex et al., 2014) of filter subsets for TN SNV (a+c) and INDEL (b+d) calls. Filters are described in Table S5. Subplots a and b show the top 25 upsets while subplots c and d show only calls that were filtered by one single filter. The plots reveal that np2's allele quality filter (AQ1) is the single most effective filter for suppressing FP SNV calls followed by strand bias (SB) and homopolymer (HP) filters. INDEL FPs are predominately filtered due to low allele frequency (AF) and also SB and HP filters. Overall, these plots were created from more than 200k individual calls in 40 different samples.

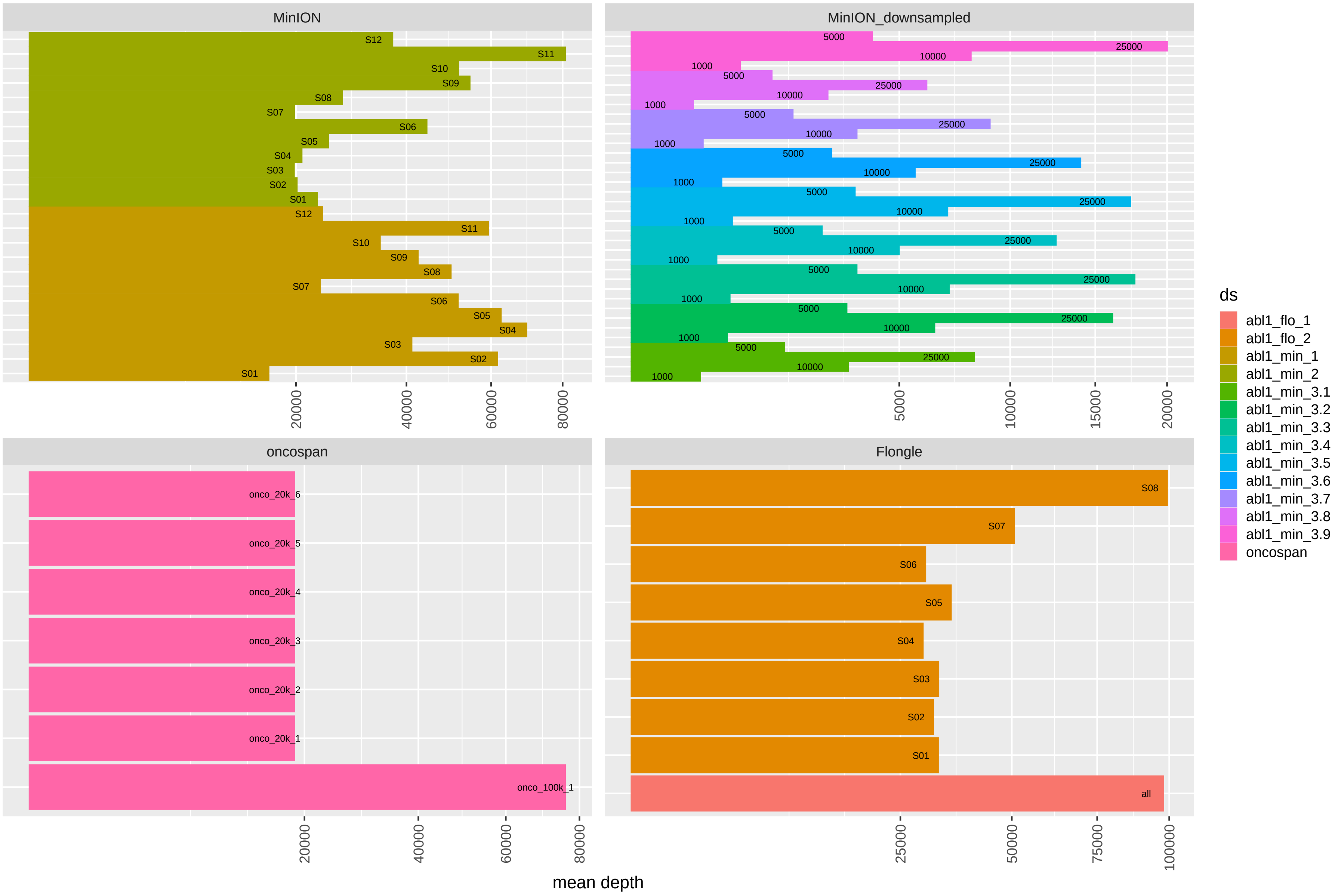

Figure S3: Mapped reads per sample, averaged over all three mappers (mm2, ngm and last) and over all sequenced amplicons.

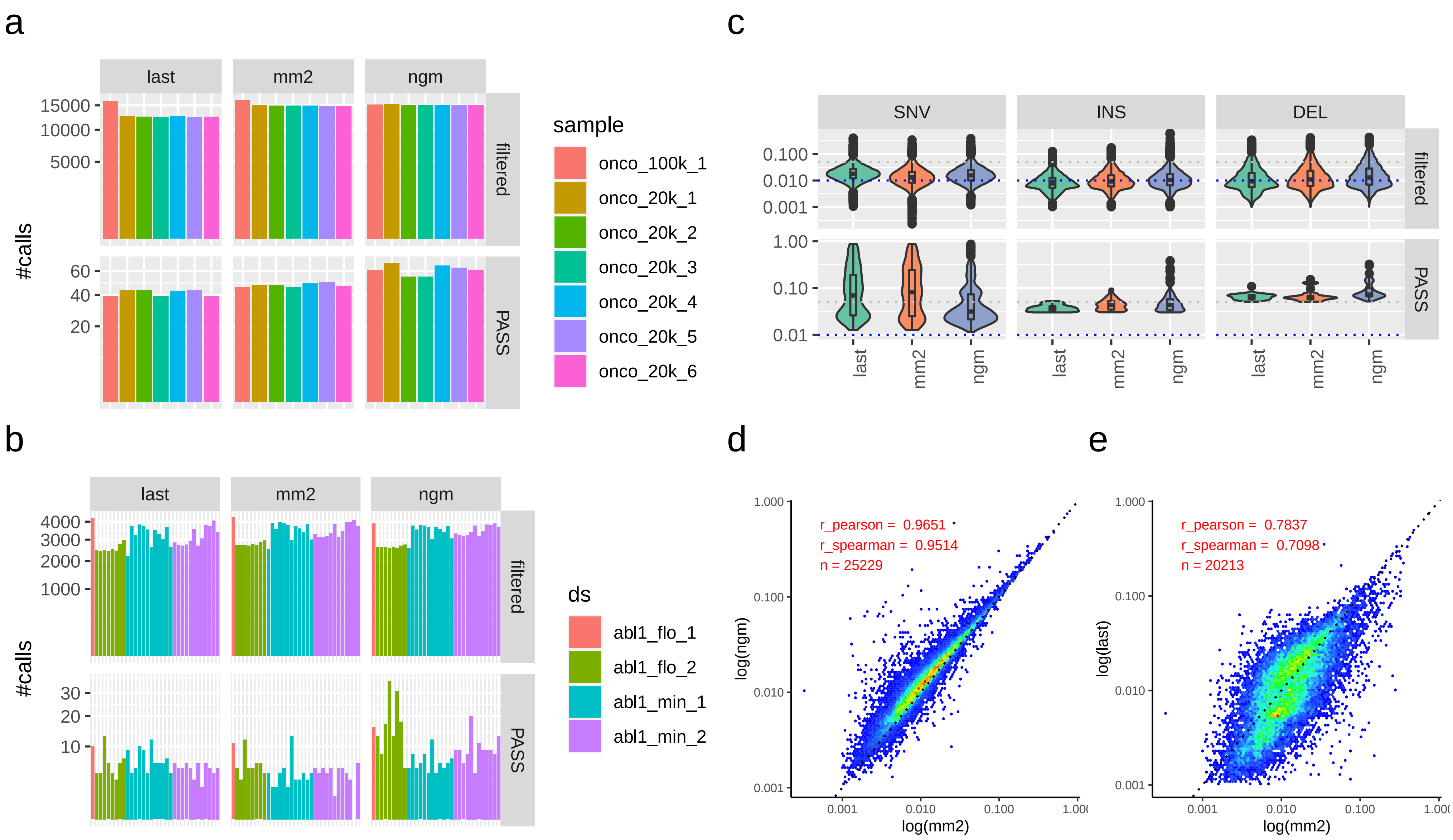

Figure S4: Subfigures a and b show the numbers of filtered and PASS calls per dataset for oncospan and abl1 data respectively and demonstrate increased numbers of unfiltered (potentially FP) calls resulting from ngm alignments. Subfigure c confirms this finding by showing slightly increased INDEL VAF distributions for ngm vs mm2 alignments. Subfigures d and e show good VAF correlation between ngm and mm2 and lower correlation between mm2 and last.

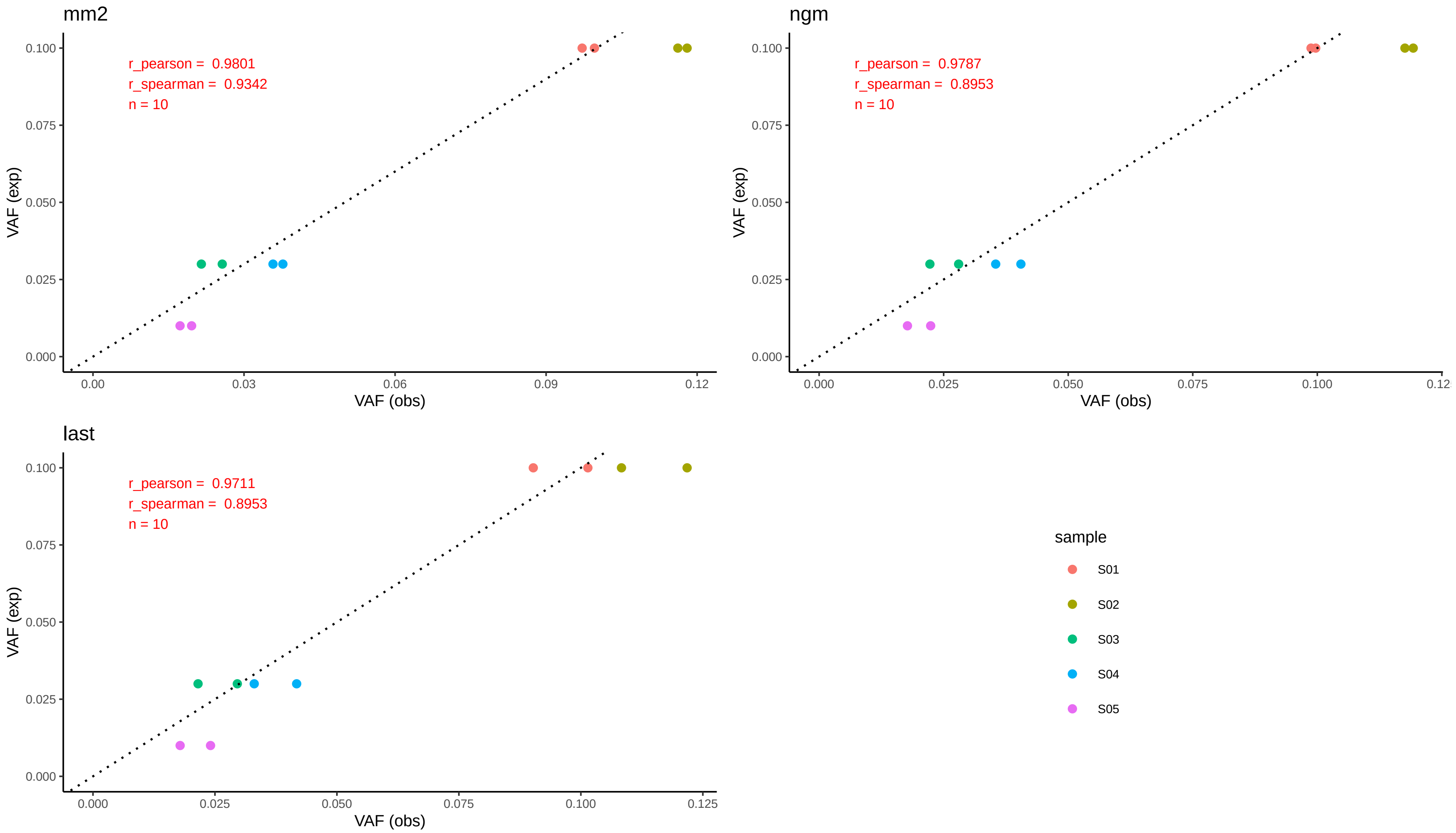

Figure S5: Correlation between expected (exp) and observed (obs) VAF for different mappers in a dilution series (abl1_min_2/dilution; Sup. Table S8) with two replicates and two target variants (E255V + T315I, compound on the same plasmid) with expected 10%, 3% and 1% VAF respectively. Sample S06 (VAF=1%) is missing as none of the respective variants was called by np2. The graph shows the best reproducibility and highest correlation between expected and observed VAF for the minimap2 alignments. It also shows very similar reported VAFs for variants linked on the same plasmid.

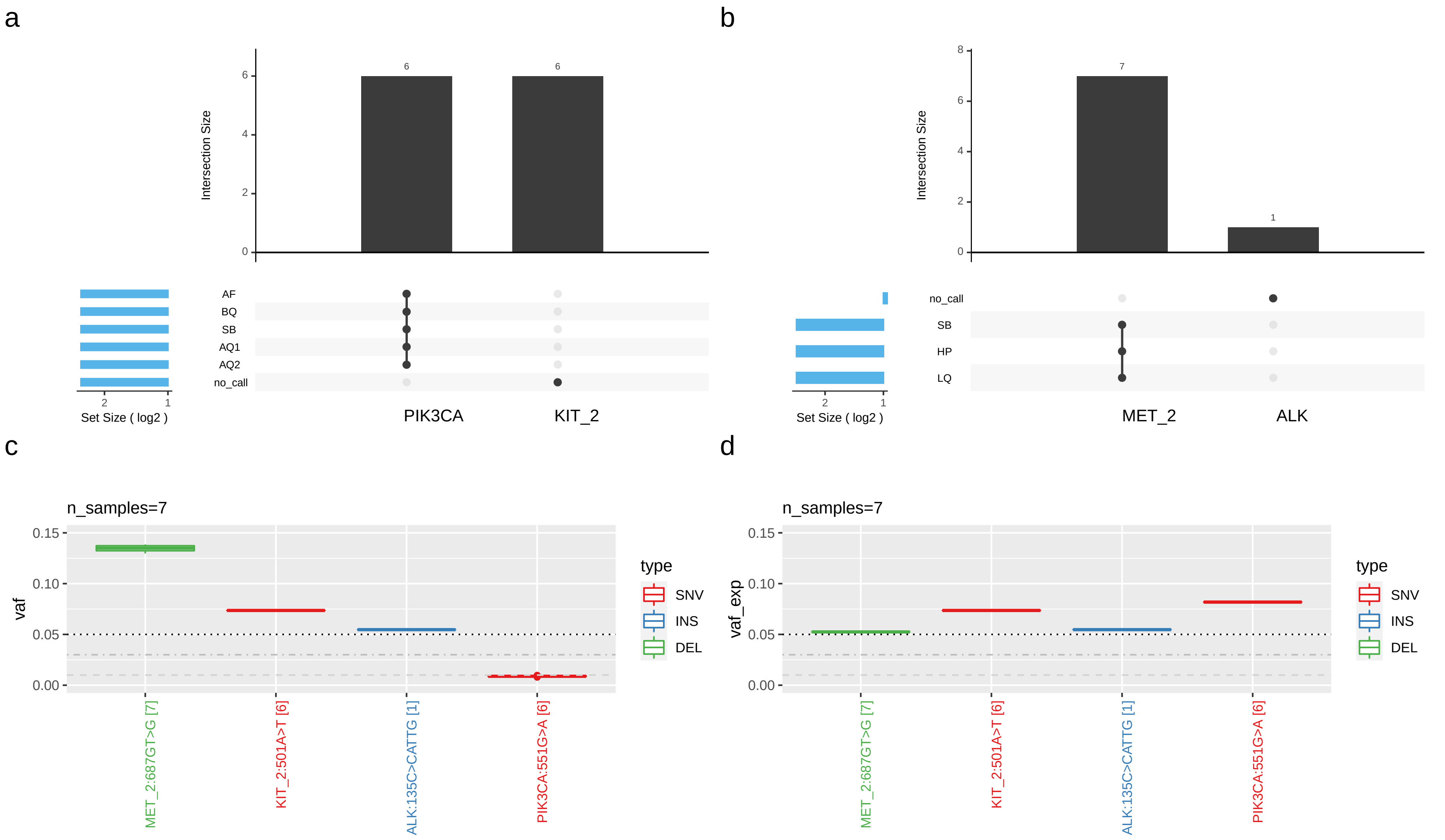

Figure S6: FN calls in the oncospan panel (7 replicates). Overall, there were 4 true variants (2 SNVs and 2 INDELs) that were filtered or not called in one or multiple replicates. Subplots a and b show the respective reasons (filters) and the numbers of replicates containing these FNs (number of top of bars). Subplots c and d show observed and expected VAF for these variants. See Sup. Table S6 for a detailed description of these FN calls.

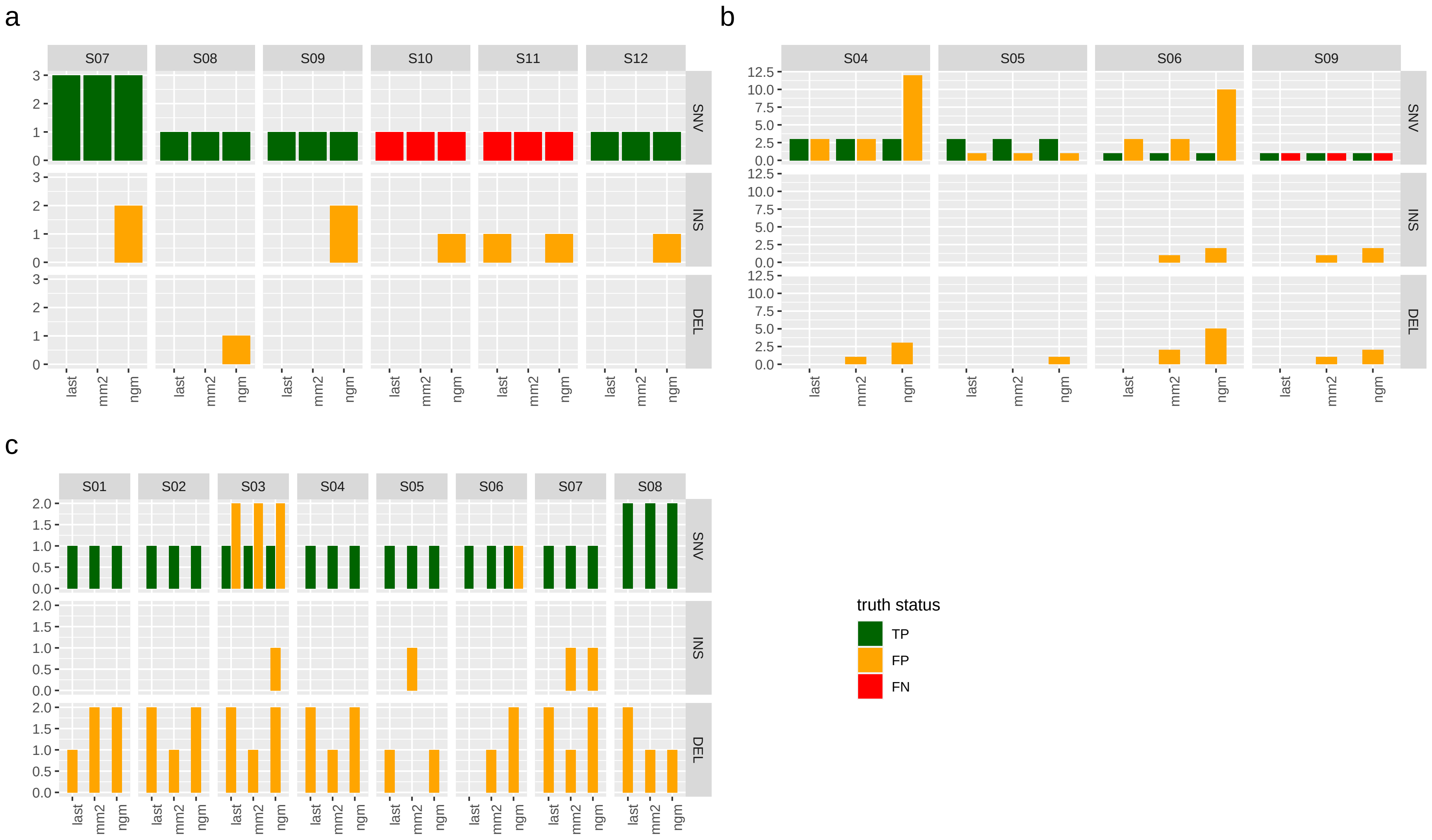

Figure S7: Detailed performance charts showing TP, FP and FN counts for all clinical samples with a known truth set. Subfigure a shows 6 ring trial samples from flowcell abl1_min_2. Subfigure b shows 4 clinical samples from flowcell abl1_min_3 (each sample was downsampled to 25k reads). Subfigure c shows 8 clinical samples from flongle abl1_flo_2. Only calls with a VAF $>$ 5% were considered in these plots. Note, that we do not have a complete ground truth for these samples due to limitations of the used validation assays (see main manuscript) which means that some of the called false positives could be true variants. Also, most false-positives have low VAF as demonstrated in Sup. Fig. S8 and Fig. 1d of the main manuscript.

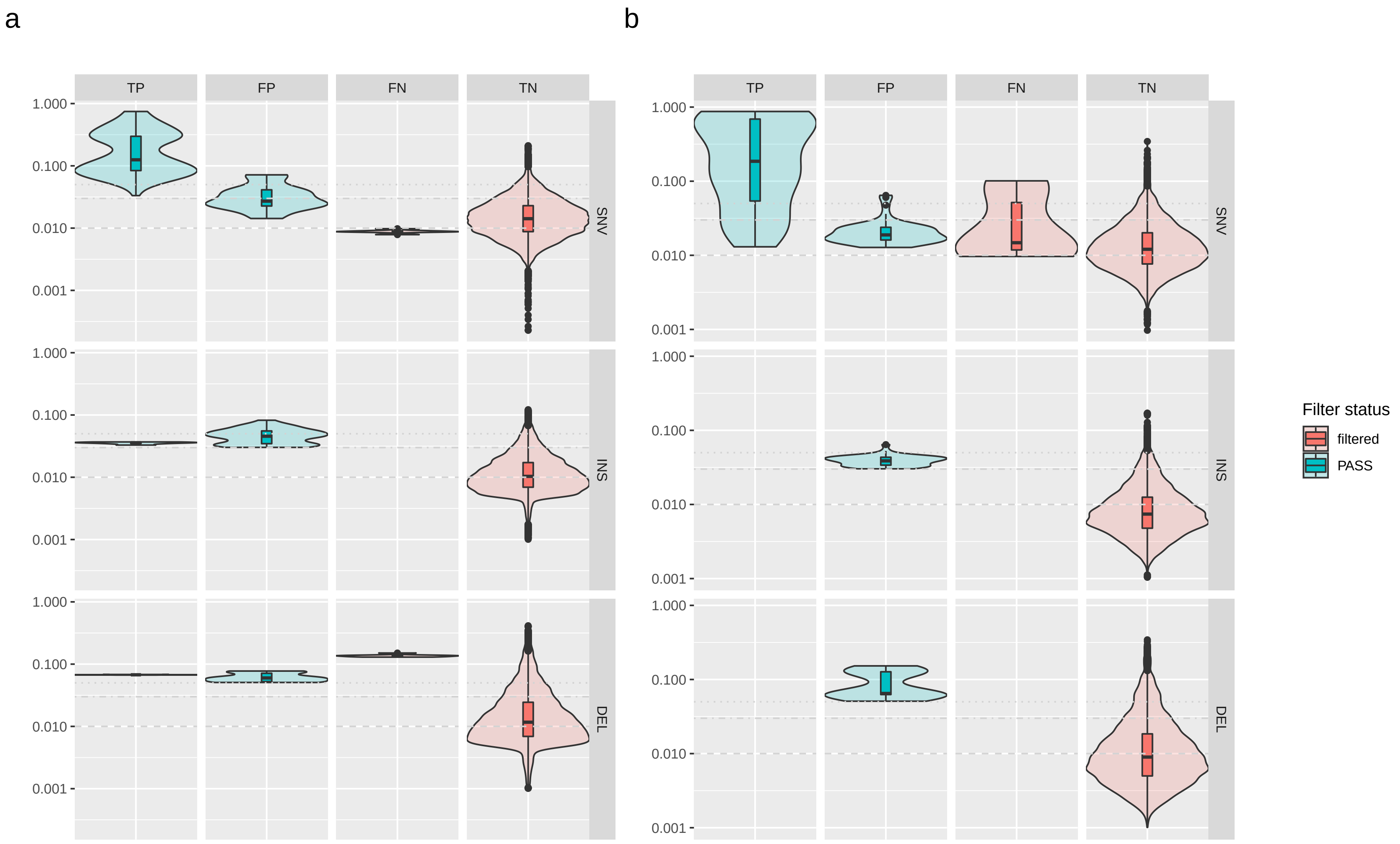

Figure S8: Distribution of observed raw VAF (y-axes; log-transformed) for filtered and unfiltered ('PASS') variants per truth status and variant type for oncospan (a) and abl1 data (b). Horizontal grey lines show calling thresholds for deletions, insertions and SNVs respectively. The graph demonstrates that np2 produces very few high-frequency FP calls despite the strong overlap of VAF distributions of filtered and unfiltered calls.

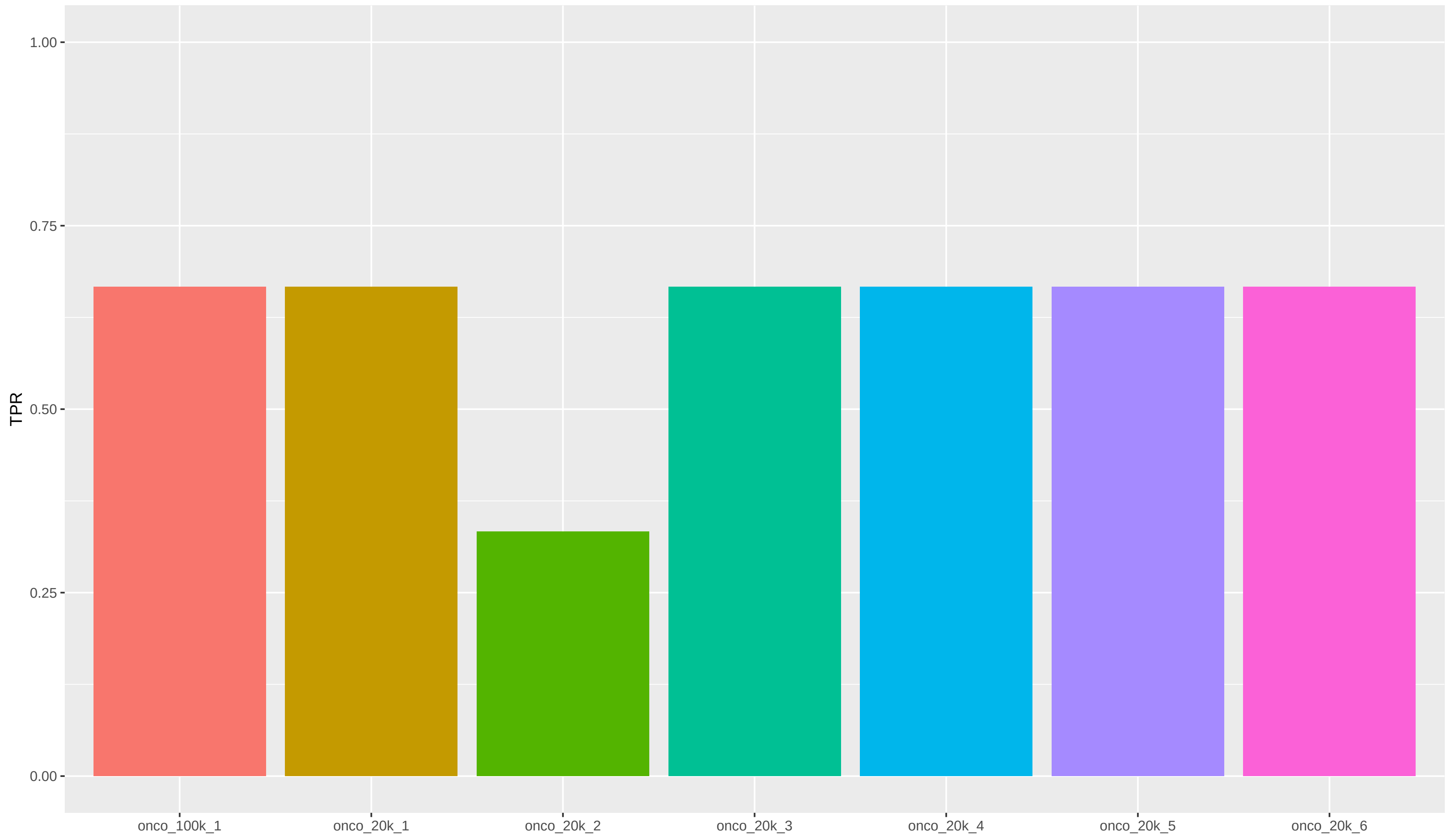

Figure S9: True positive rate for three validated INDEL calls over all oncospan replicates.

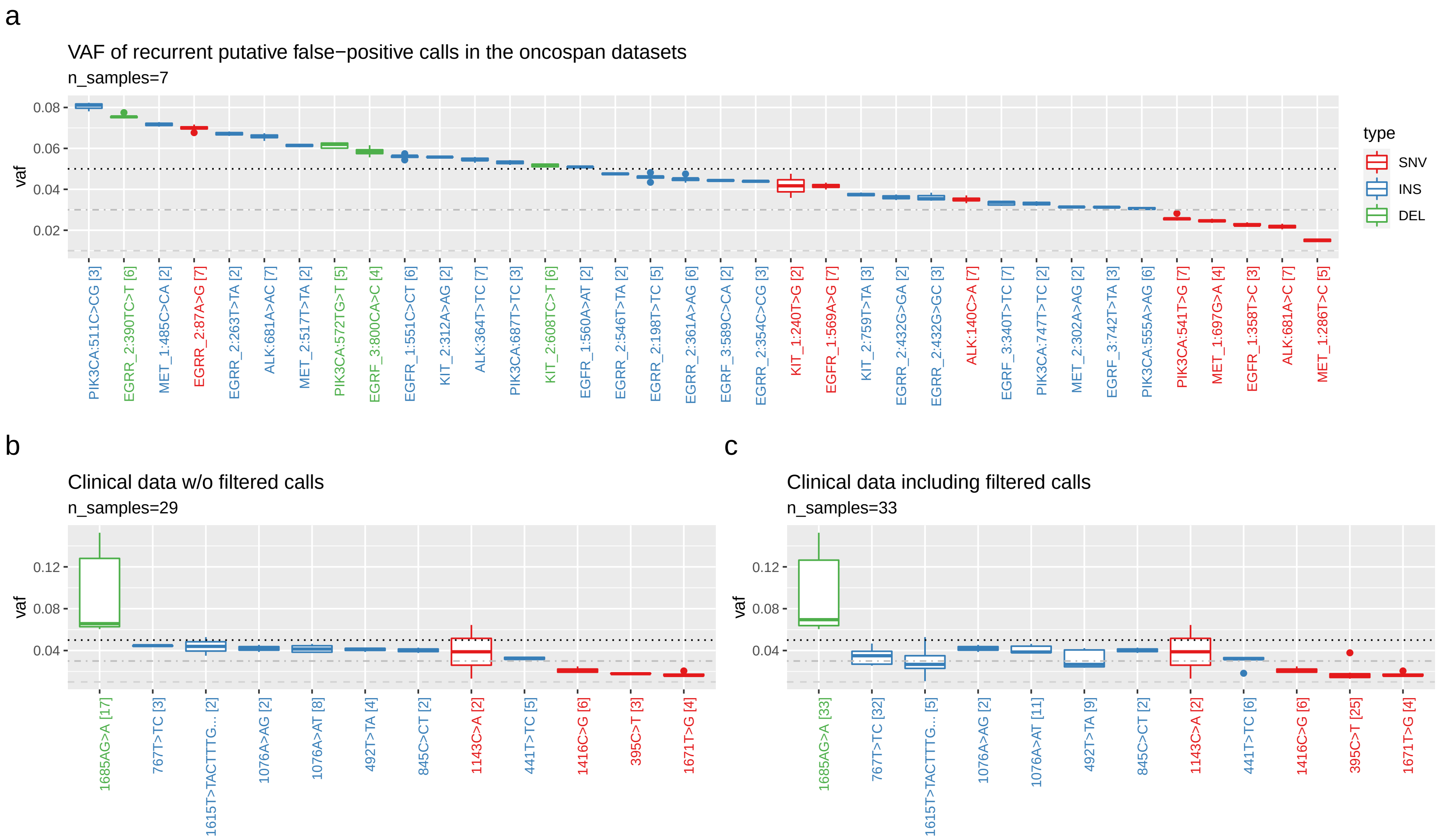

Figure S10: VAF distributions of putative false-positive calls in the oncospan data (a) and all clinical datasets (b) that are found in more than one sample. The boxplots contain all observed VAFs from all considered datasets and boxes are colored by variant type. Dotted, dotdashed and dashed lines indicate the calling thresholds for deletions, insertions and SNVs respectively. Numbers in square brackets indicate the number of datasets a FP was found in (in subfigure c this includes filtered calls). Only one sample per patient was considered for subplots b and c to not artificially inflate the statistics by replicated datasets. Subplot a shows that there are no high-VAF oncospan false positives (max VAF ∼8%), that most recurrent FP calls are small insertions and that observed allele frequencies are very similar across the 7 replicates (narrow boxes). Subplot b and c confirm that most recurrent FP calls are small, low-frequency INDELs and show one highly recurrent, possibly systematic FP at 1685AG>A which we consider a PCR artefact (called but partially filtered in all 33 considered abl1 samples). Note that some of the reported FP calls might actually be true but could not validated due to the detection limits of the used orthogonal technology (e.g., ∼20% VAF for Sanger sequencing).

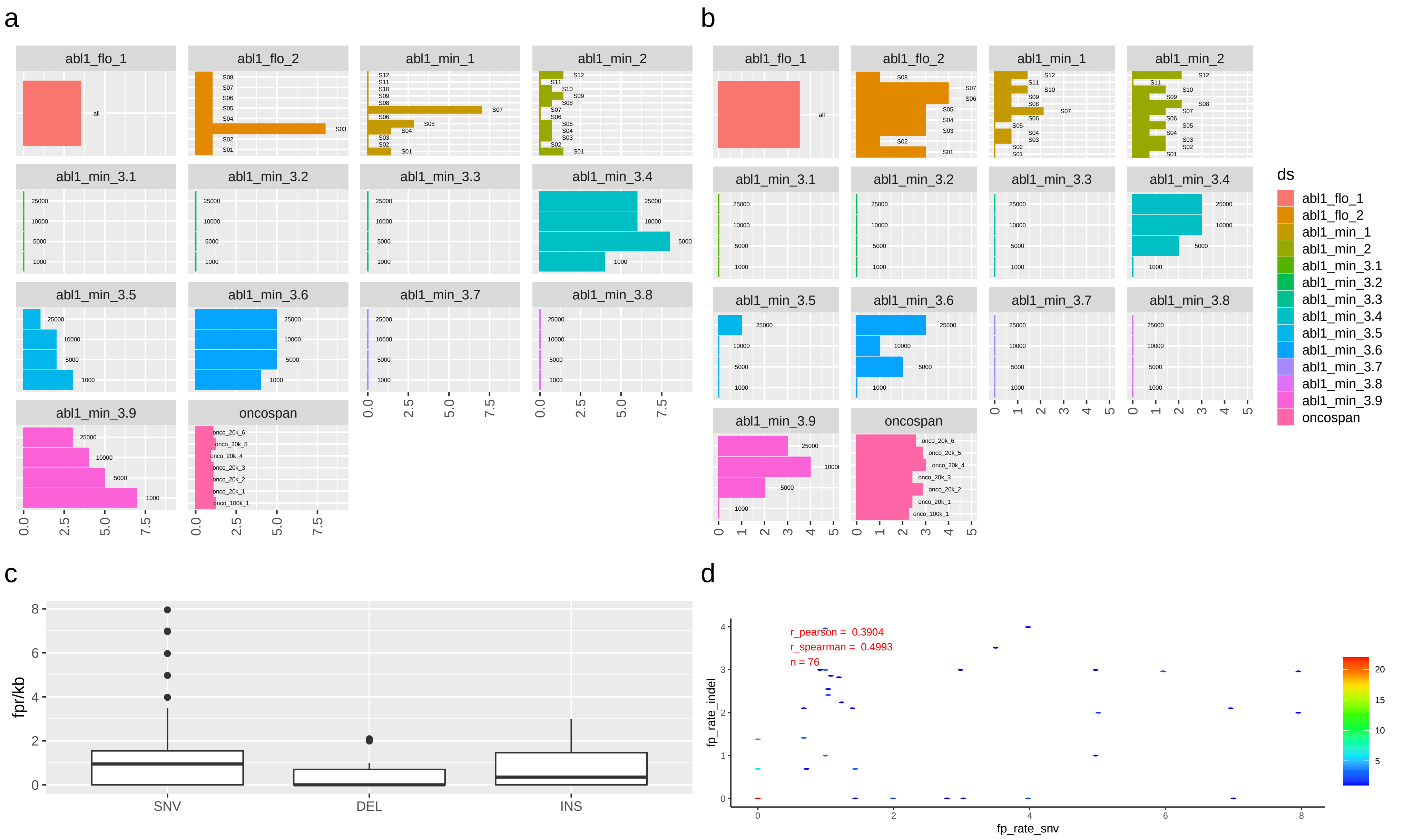

Figure S11: False-positive rates (FPR) per kb per sample for SNVs (a), INDELs (b) as well as overall rates per variant type (c). Note, that for some abl1_min3 downsamples we observed smaller FPR in samples with lower read counts. Overall, we found median FPR values of 0.95 and 0.7 for SNVs and INDELs respectively. Subfigure d shows that SNV and INDEL FPR values are weakly correlated.

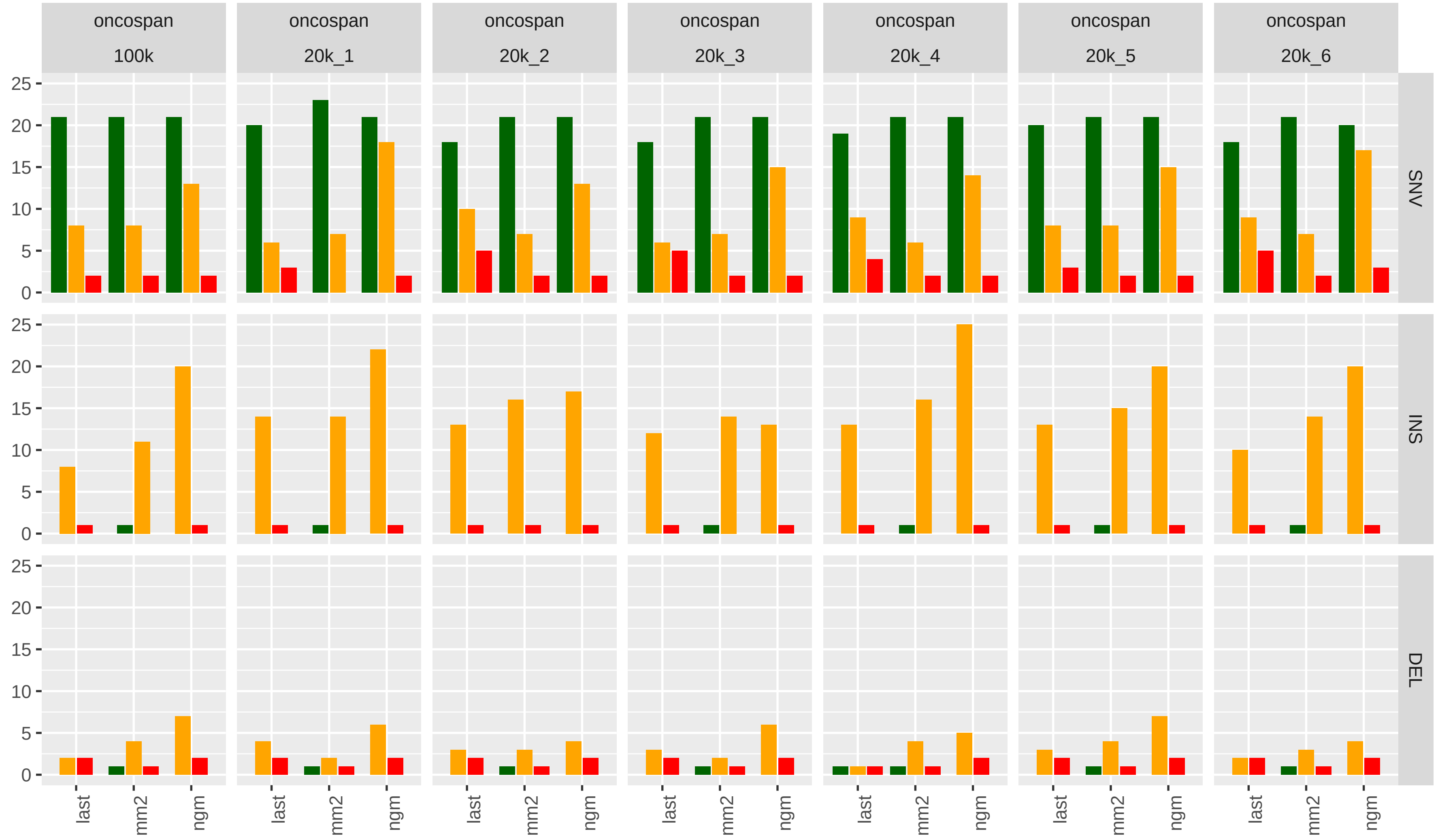

Figure S12: Number of true positive (TP), true negative (TN) and false negative (FN) variant calls per oncospan dataset, caller and variant type for all calls with an observed or expected (for FN) VAF $> 1\%$. Note that most of the shown FP have low VAF, cf. Fig. 1 in main manuscript.

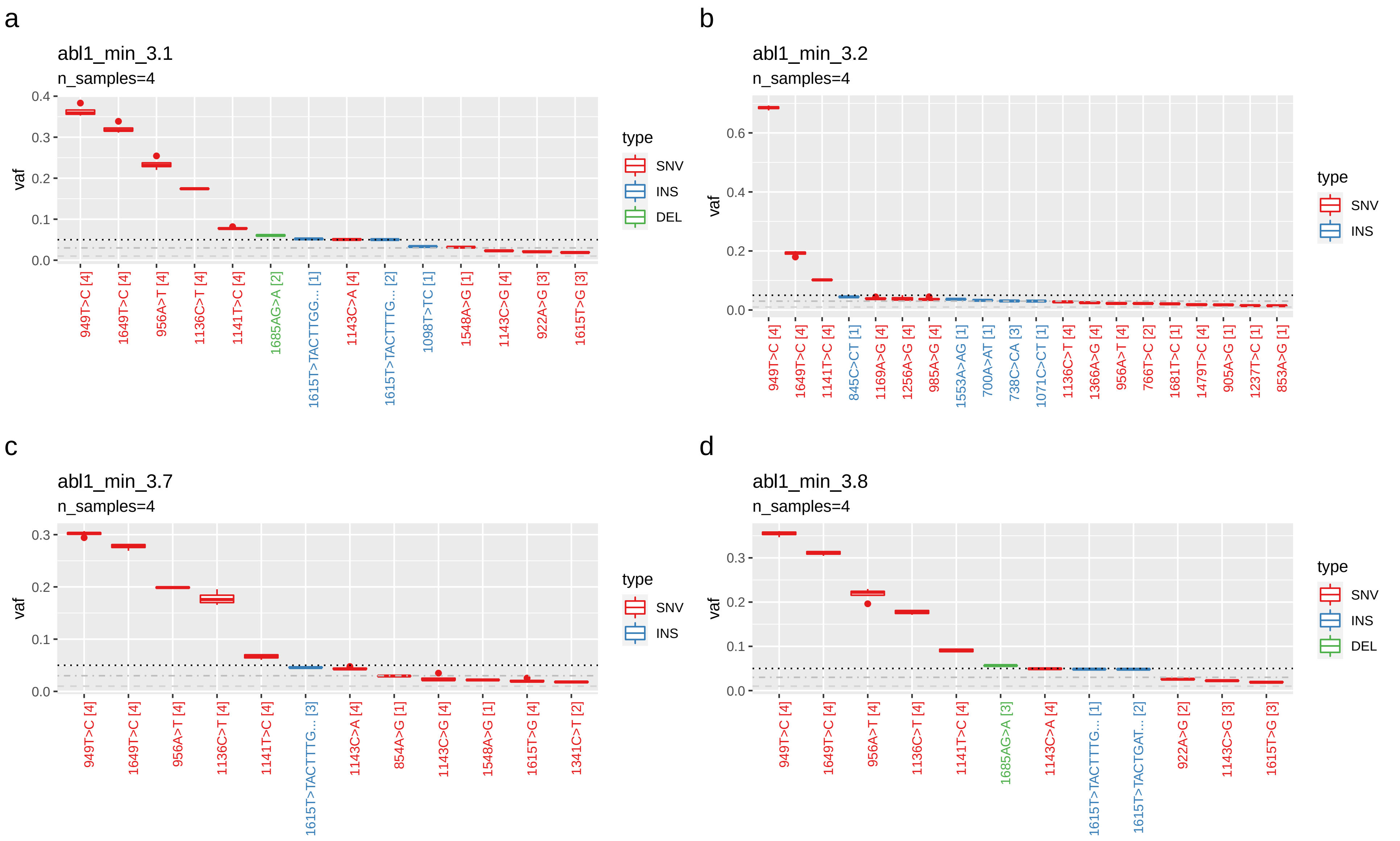

Figure S13: Variant calls, their VAF and their recurrence across 4 downsamples (1k, 5k, 10k and 25k reads) in four exemplary abl1 samples from two patients (a+b: patient1, c+d: patient2). Variants are sorted by mean VAF, the number of downsamples a variant was called in is given in square bracket, i.e. a value of 4 corresponds to calls found in all four downsamples. Together, these boxplots show that the estimated variant calls and VAFs are highly reproducible across the downsamples despite the coverage differences and the low coverage of the 1k reads subsample.

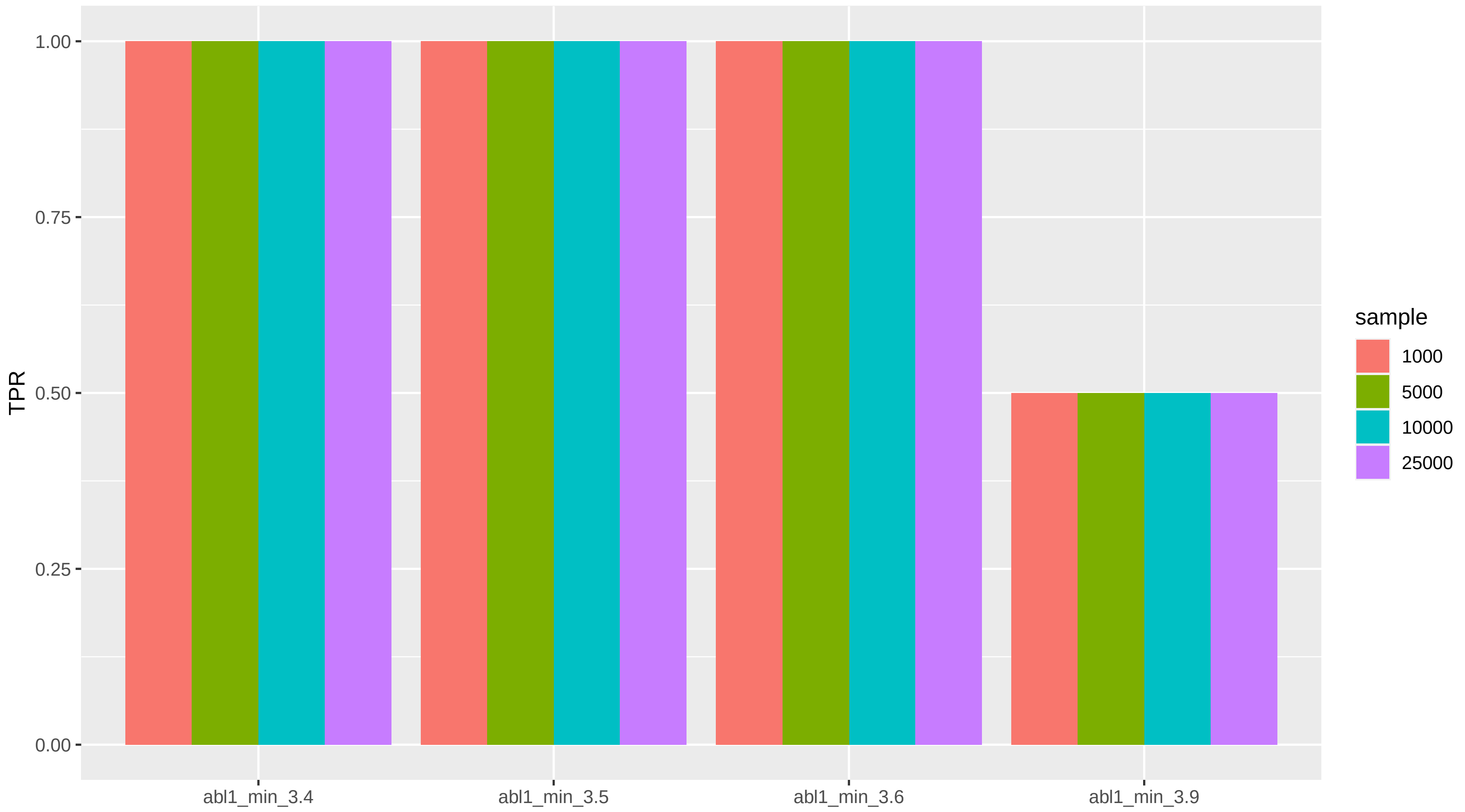

Figure S14: True positive rate (TPR) for all downsamples (1k, 5k, 10k and 25k reads) of 4 clinical samples from flowcell abl1_min_3 with an existing truth set. All but one variant in sample 'abl1_min_3.9' were found in all downsamples. This FN is E255K which is always filtered by our 'HP' filter, see discussion in Methods section of main manuscript.

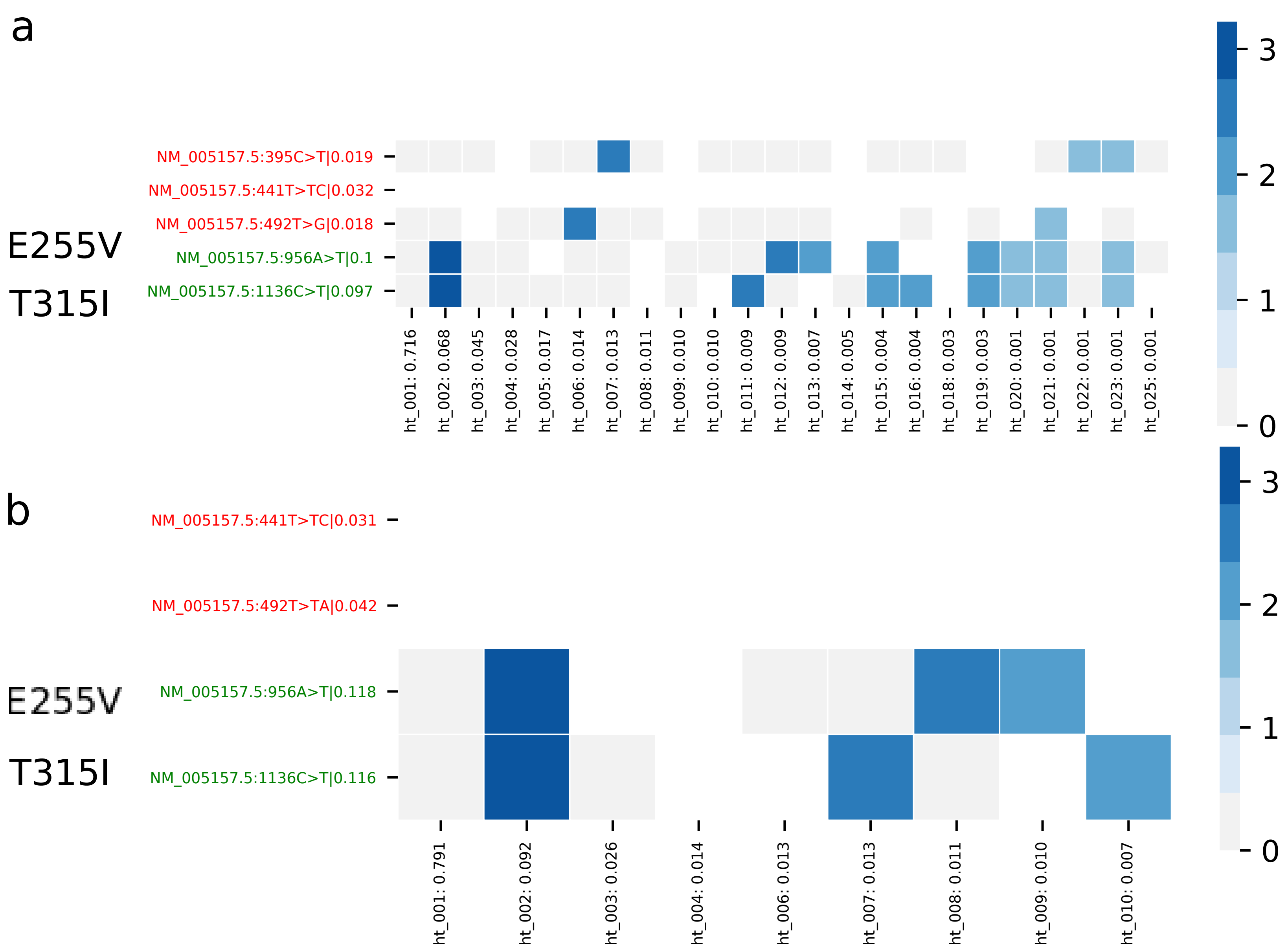

Figure S15: Haplotype map for samples 1 (a) and 2 (b) of flowcell abl1_min_2. Expected VAF for the compound mutations E255V and T315I was 10% for both samples. Red mutations are false-positives, see caption of main Fig. 2 for a detailed description of our haplotype map format.

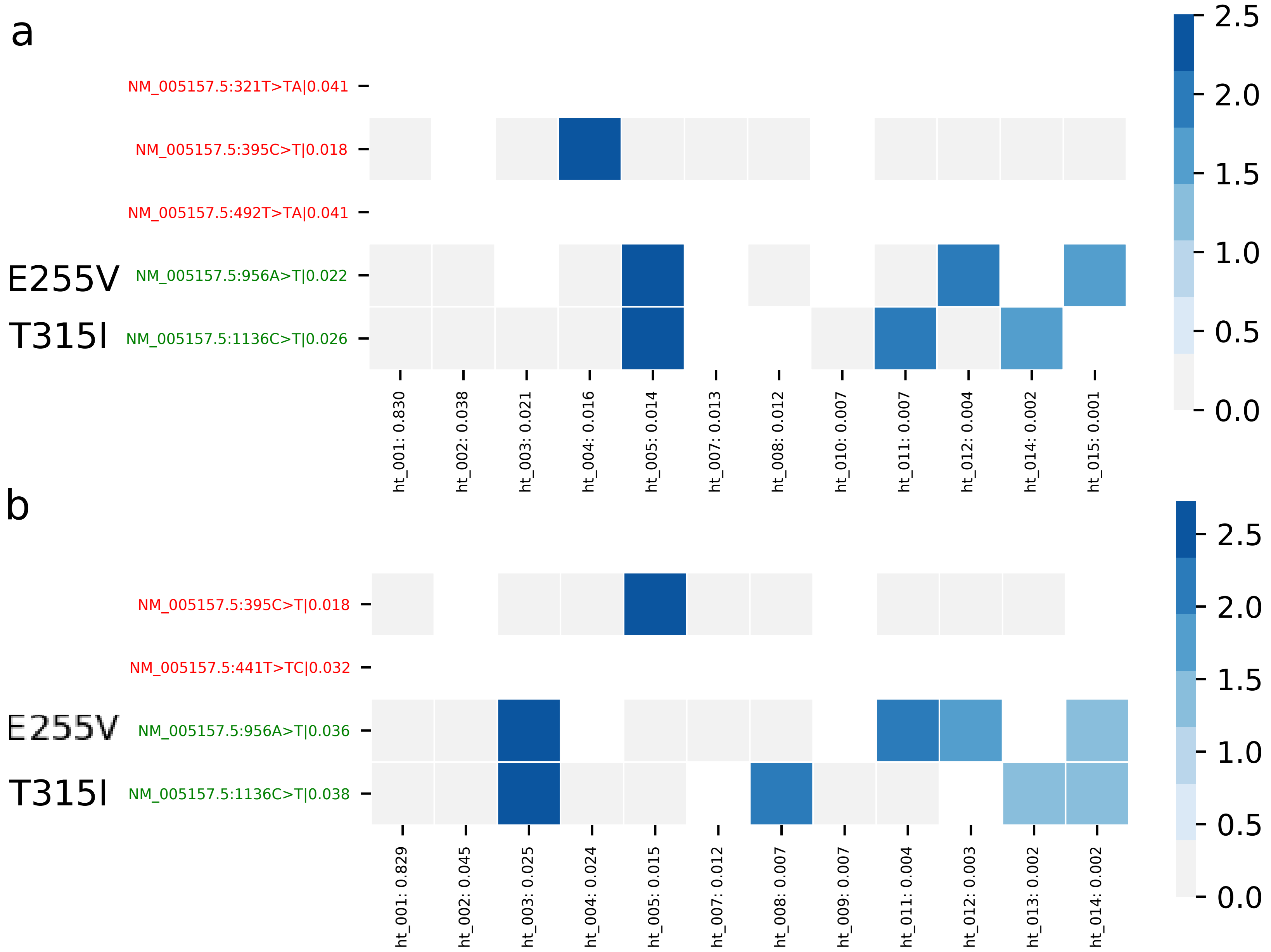

Figure S16: Haplotype map for samples 3 (a) and 4 (b) of flowcell abl1_min_2. Expected VAF for the compound mutations E255V and T315I was 3% for both samples. Red mutations are false-positives, see caption of main Fig. 2 for a detailed description of our haplotype map format.

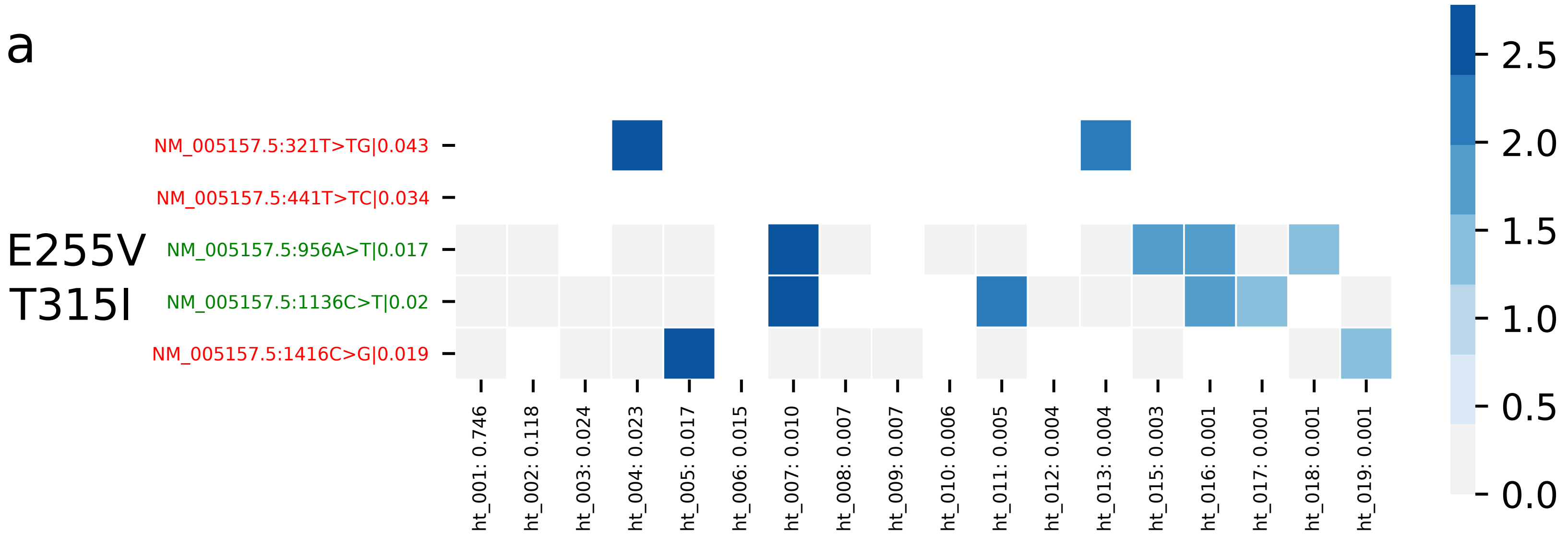

Figure S17: Haplotype map for sample 5 of flowcell abl1_min_2. Expected VAF for the compound mutations E255V and T315I was 1%. Sample 6 (a replicate of sample 5) is not shown as np2 called none of the two TP mutations in this replicate. Red mutations are false-positives, see caption of main Fig. 2 for a detailed description of our haplotype map format.

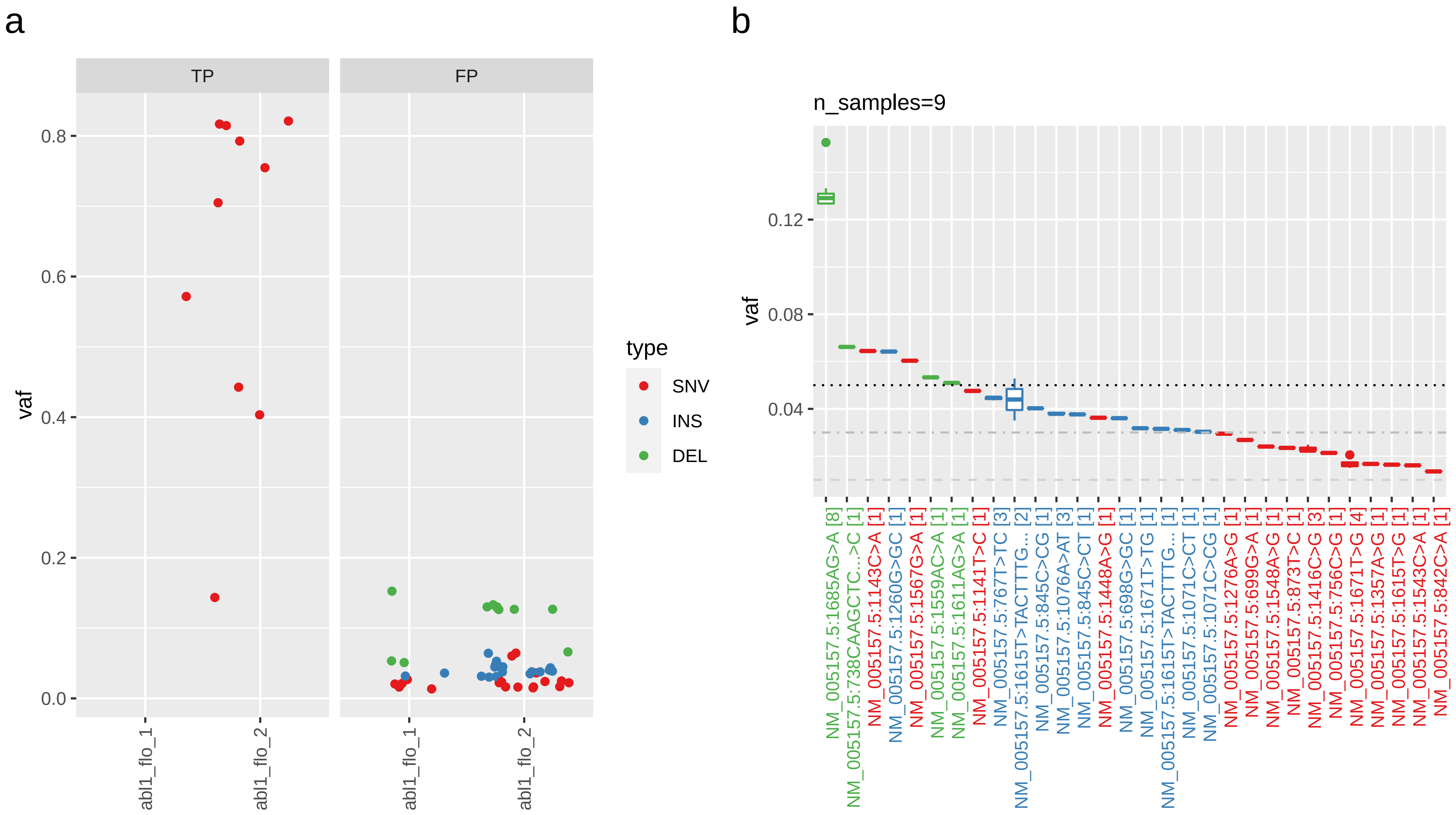

Figure S18: Subfigure a plots VAF of TP and potential FP calls in the flongle datasets (abl1_flo_1: 1 benchmark sample, abl1_flo_2: 8 clinical samples from 8 different patients). Subfigure b plots recurrence of FP calls (number of samples containing a variant given in square brackets; cf. Sup. Fig S10 for a detailed explanation of these plots) and shows the same deletion at 1685AG>A as found in all other abl1 samples which we consider a PCR artefact (cf. Sup. Fig S10 b + c). All other recurrent FPs are low VAF insertions and SNVs. Note that we don't know whether all low-frequency FP variants are actually false-positives due to the detection limits of the used validation methods.

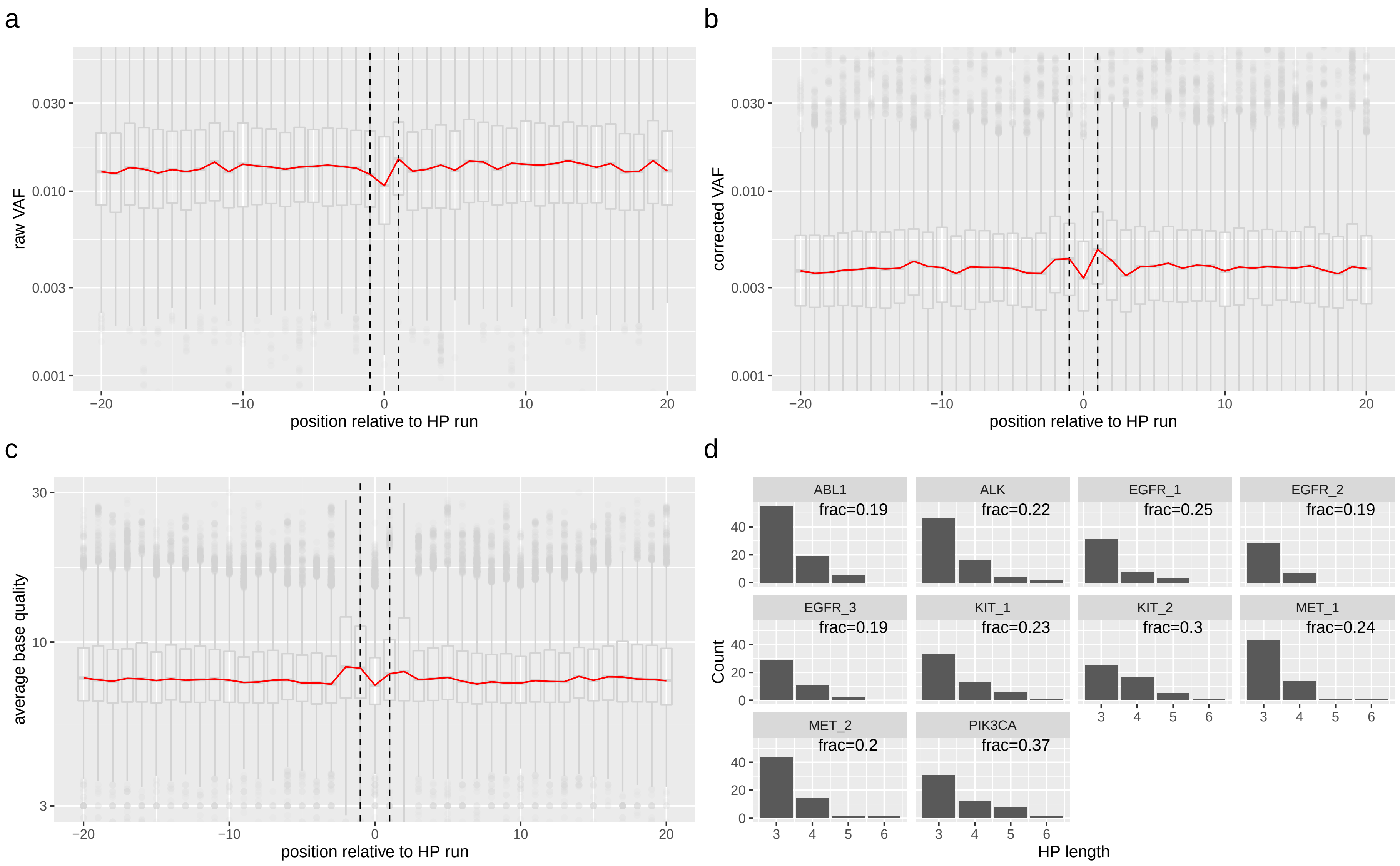

Figure S19: Subfigures a and b show median raw and corrected VAF (red line) around homopolymer runs with a minimum length of 3. The X-axis shows genomic positions relative to start/end of hp runs where -1/+1 refer to the first/last base of the hp run respectively (dashed black lines) and all bases in the ”body” of the hp run are averaged and summarised at position zero. The figure shows that our correction algorithm filters around two-thirds of base calls and overall results in relatively increased VAF around hp run borders. Note that these positions also show increased average base qualities (c). Subfigure d shows a histogram of all hp runs per amplicon. Labels in the plots show the fraction of amplicon bases in the respective hp runs. Subfigures a-c were calculated from 40 samples containing data of over 5500 hp runs.

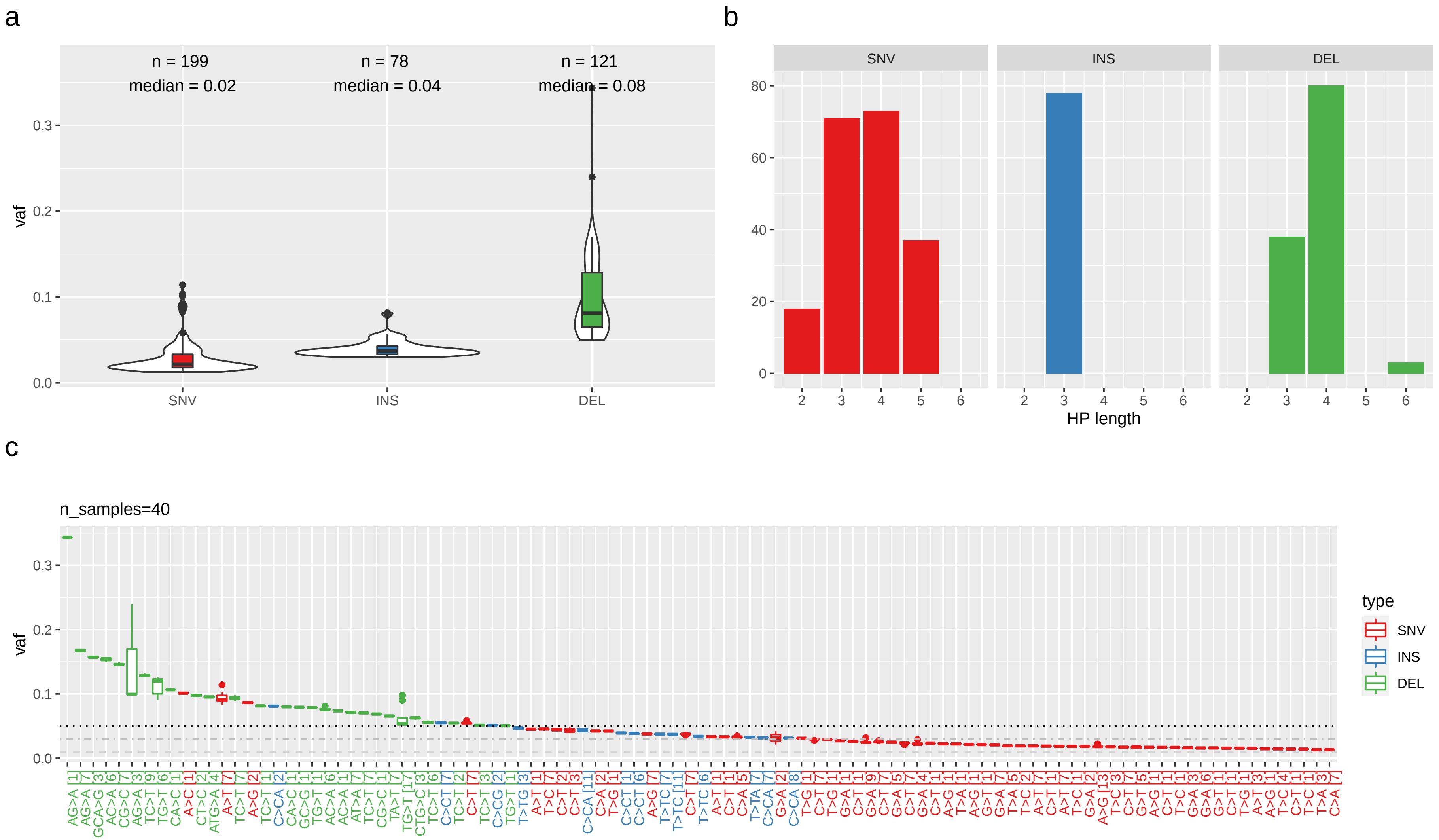

Figure S20: VAF, hp length and recurrence statistics for all calls from 40 samples that were filtered exclusively by our HP filter. Only 1 of 398 calls (0.25%) was a FN, all other were true negatives. Subfigures a and c show that we found mostly low-frequency calls with a high degree of recurrence (99 distinct calls; one deletion was, e.g., called in 17/40 samples) suggesting systematic reasons for such calls that could be exploited for improved filtering. Subfigure b shows a histogram of hp lengths in the context of such calls. For SNVs, the plotted values correspond to the maximum length of an up/downstream hp run). For INDELs, the length is the sum of up- and downstream hp runs of the first base of the respective alt allele (see Methods in main manuscript).

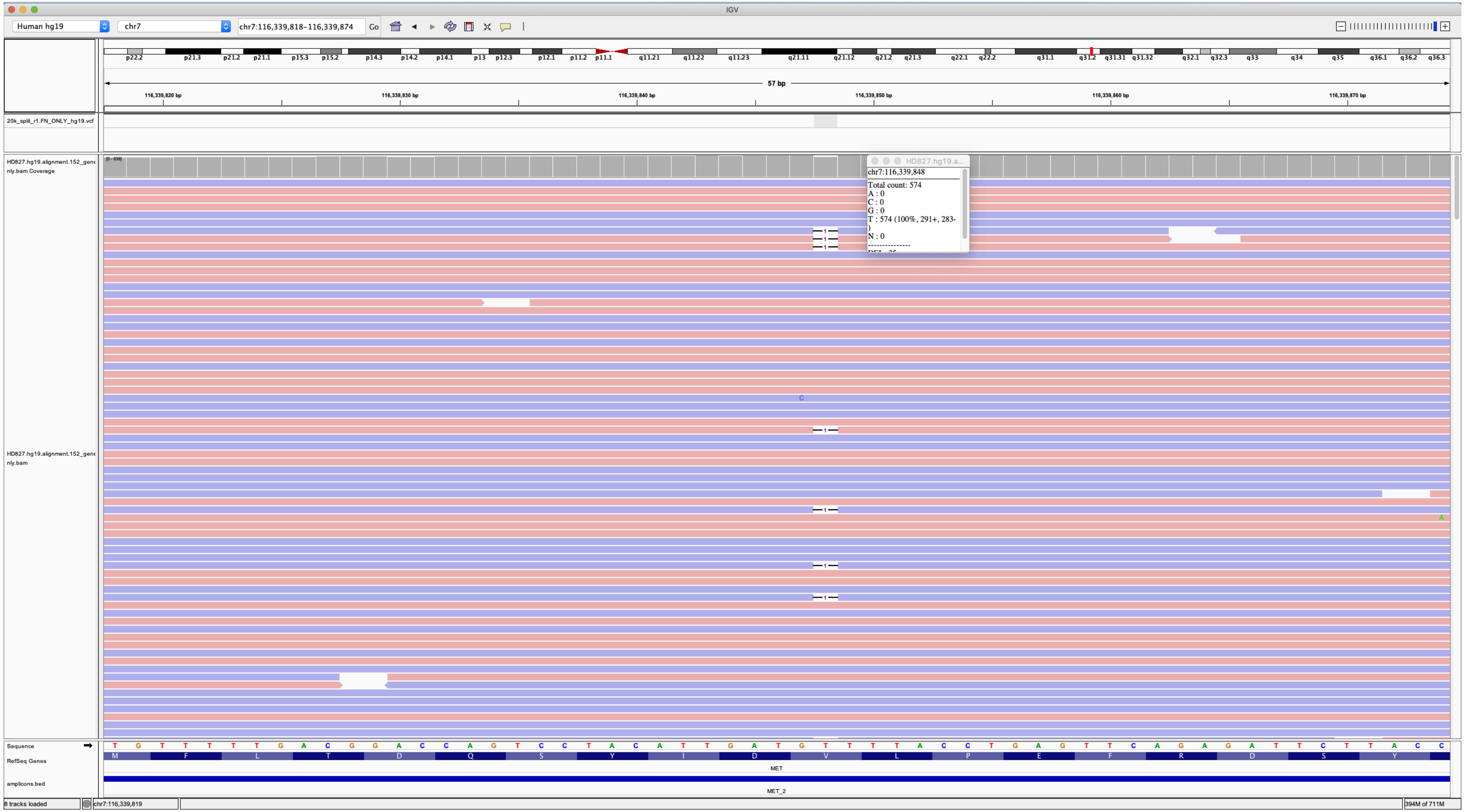

Figure S21: IGV screenshot of the oncospan WES data showing a 1bp deletion at MET_2:687GT>G. The deletion is clearly visible and was called by pisces in the deep short read alignment (read depth: 572, VAF=0.05245) but filtered by np2 resulting in a FN call, cf. Sup. Table S6.

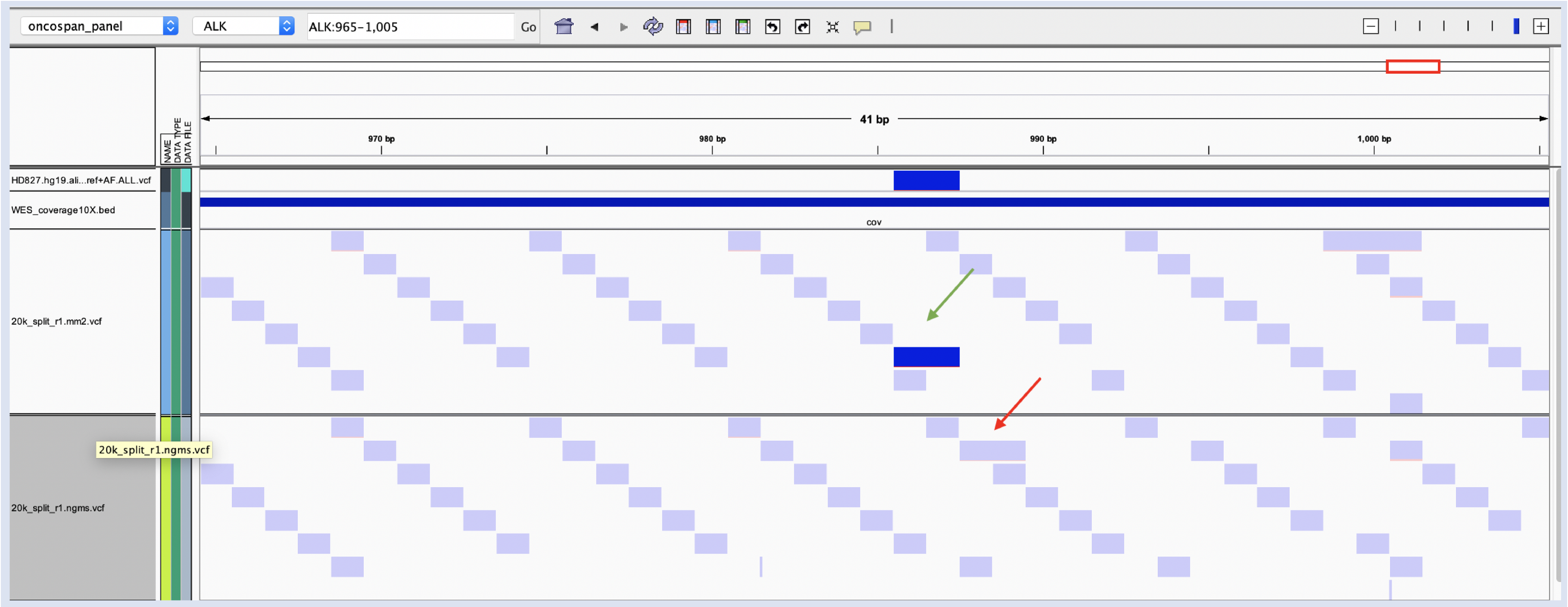

Figure S22: IGV screenshot demonstrating local differences in read alignment that result in differing overall performance of the respective mappers. Shown tracks (from top) are: (i) pisces calls from the WES data (truthset) in the ALK amplicon of an oncospan 20k replicate, (ii) sufficiently covered regions in WES data, (iii) np2 calls resulting from mm2 (top) and ngm (bottom) alignments. Filtered and unfiltered calls are shown as light-blue and dark-blue rectangles respectively. The screenshot shows a single true-positive INDEL call in the mm2 alignment (green arrow) while the same INDEL is shifted to the right in ngm due to its different alignment algorithm and filtered by np2 (red arrow).

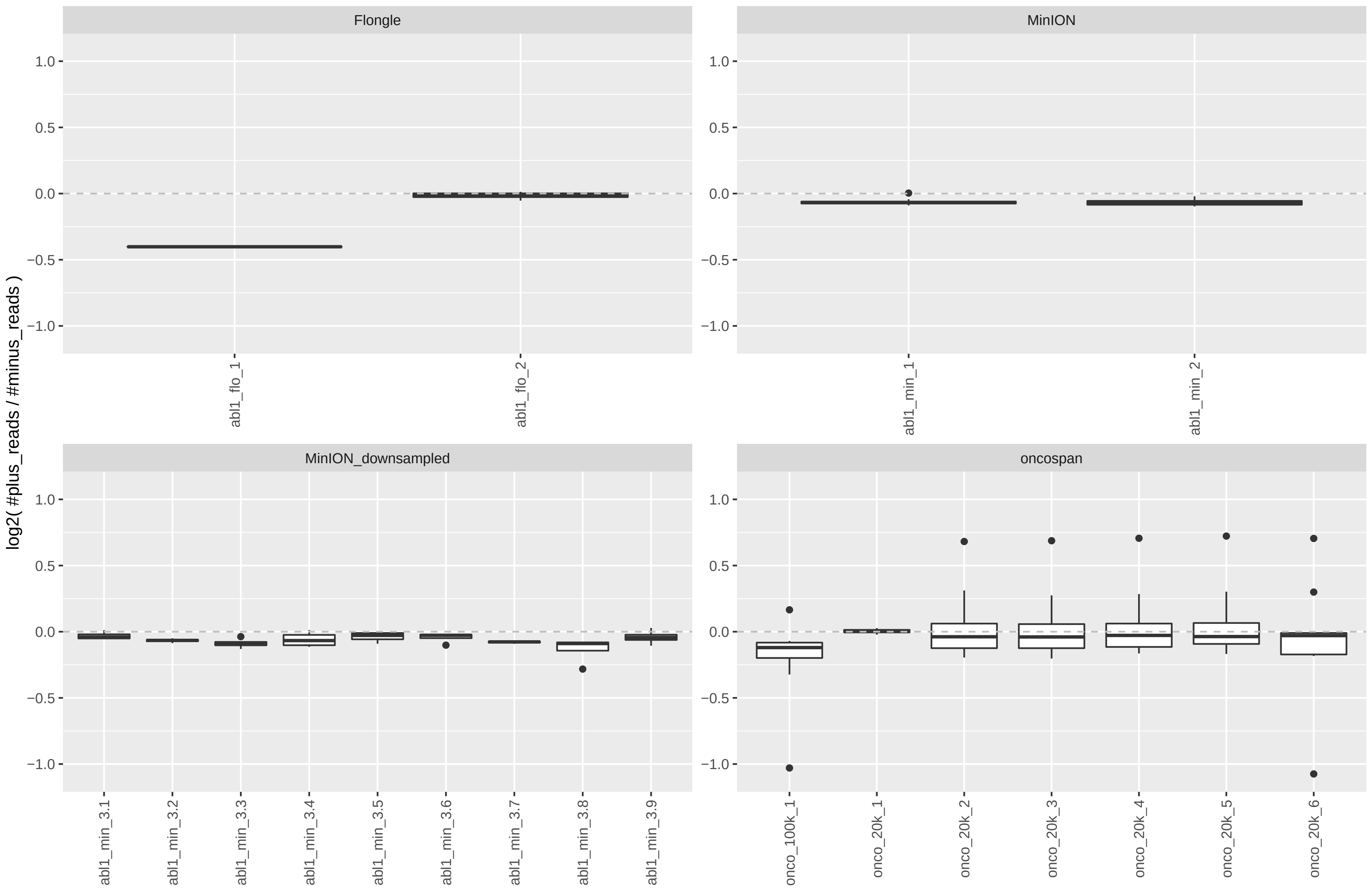

Figure S23: Imbalance of read counts per strand for all datasets/samples: log2(#plus_strand_reads / #minus_strand_reads). Please note that nanopanel2 corrects for this sample-wide imbalance to avoid over-filtering due to strand bias (see Methods in main manuscript).
